# Polyamine biosynthesis dysregulation in Alzheimer’s disease and Down syndrome cellular models

**DOI:** 10.1101/2025.01.31.635912

**Authors:** Andres Sola, Alex Sandberg, Caitlin Pham, Alexandra Revier, Mia Hebinck, Alexandra Penney, Pablo Caviedes, Sunil Kumar, Ann-Charlotte Granholm, Daniel A. Linseman, Daniel A. Paredes

## Abstract

**BACKGROUND:** Individuals with Down Syndrome (DS) frequently develop early onset Alzheimer’s disease (AD) with pathological hallmarks closely resembling AD due to several triplicated genes on chromosome 21. Polyamines are small, organic molecules that play a pivotal role for growth and differentiation, and a dysregulation of polyamine pathways is implicated in AD pathology. However, their role in DS-associated AD is unclear.

**METHODS:** We analyzed polyamines and their metabolite levels in mouse hippocampal cells and human DS-AD and AD hippocampal tissue and assessed the effects of the ODC inhibitor difluoromethylornithine (DFMO) on Aβ42 aggregation and protein expression in DS fibroblasts.

**RESULTS:** Amyloid-β42 increased polyamine levels via ornithine decarboxylase (ODC) activation in a dose-dependent manner. DFMO reduced Aβ42 aggregation, decreased amyloid precursor protein (APP) levels, and normalized proteins linked to AD pathology in DS fibroblasts. Polyamine levels were elevated in DS-AD hippocampal tissue, with colocalization of ODC and Aβ42 aggregates.

**CONCLUSION:** These findings suggest that polyamine biosynthesis may exacerbate Aβ42 toxicity and APP expression, contributing to AD progression in DS. The ability of DFMO to reduce Aβ42 aggregation and restore protein homeostasis presents the polyamine pathway as a therapeutic target for DS-AD management.

## BACKGROUND

Down syndrome (DS) and Alzheimer’s Disease (AD) share underlying pathological mechanisms that often lead to progressive dementia [1–3]. The amyloid precursor protein (*APP*) gene, a known risk factor for AD, is triplicated in DS, and is located on human autosome 21 [4]. This results in high levels of toxic cleavage products including amyloid-beta (Aβ), with accumulation of amyloid plaques typically beginning in DS after 30 years of age. Other genes on Chr. 21, including e.g. the superoxide dismutase 1 (SOD1) gene, genes for several interferon receptors, dual specificity tyrosine phosphorylation regulated kinase 1A (Dyrk1A) and regulator of calcineurin 1 (RCAN1) are known to affect AD pathology including the amyloid, phospho-tau, and neuroinflammatory and oxidative stress pathways in the brain [5,6]. The amyloid plaque load increases progressively, beginning in the frontal cortex and eventually reaching all brain regions, although diffuse plaques have been observed in the hippocampus in those with DS in their teens and 20s [7–9]. Shortly after the accumulation of the first amyloid plaques, intracellular aggregation of hyperphosphorylated tau (p-Tau) forming neurofibrillary tangles (NFTs) begins, eventually leading to a Braak stage of V-VI in most individuals with DS, along with additional co-pathologies including neuronal loss, activation of glial cells, and hippocampal atrophy [10].

Numerous hypotheses exist to explain the cumulative and widespread death of neurons in both cortical and subcortical brain regions and the emergence of amyloid plaques and NFTs that characterize AD (for review see [11]). The amyloid cascade hypothesis posits that increased APP abundance causes Aβ peptide to aggregate into plaques, inducing oxidative damage and mitochondrial dysfunction, in turn leading to cell death [12,13]. Previous work has shown that by accumulating in the mitochondrial matrix, Aβ promotes the generation of reactive oxygen species (ROS), leading to neuronal apoptosis [14–16]. However, the progression of AD-like pathology in those with DS is a complex process, involving interactions between many genetic aberrations as described above.

Polyamines (PAs) are ubiquitous small polycations involved with numerous functions in cells, including cellular proliferation, gene regulation, and autophagy through their ability to interact with negatively charged molecules [17–26]. They are also known to respond to reactive oxygen species (ROS) with increased levels which is further exacerbated by the presence of Aβ [27]. Previous animal and human studies including *post-mortem* AD brain analyses have shown increased levels of key enzymes in the PA pathway [28–31]. Furthermore, metabolomic studies show alterations in the PA pathway in both DS and AD [32,33]. An increase in PA levels, otherwise known as the “PA response,” results in enhanced PA biosynthesis with similar increases in exogenous PA uptake [34–36]. The rate-limiting step of the synthesis of PAs occurs when ornithine is processed by ornithine decarboxylase (ODC) to yield putrescine, followed by spermidine and spermine through the addition of amine groups. These small molecules are capable of regulating their own production broadly through inhibition of ODC by translational control or individually by catabolism to revert to former PAs. However, catabolism can lead to increased potential for oxidative damage by the production of the ROS hydrogen peroxide [37] and the toxic aldehyde acrolein [38–42]. The subcellular location of ODC, as well as ODC and PA levels, are dysregulated in AD brains, suggesting that alterations in the PA biosynthesis occur as a result of AD pathology, or may be involved in the accumulation of AD pathology [29,32,43].

Research findings on the role of PAs in AD pathogenesis have been paradoxical, suggesting that accumulations of PAs can be both protective and harmful to the cell via many different pathways and neuronal/glial interactions [44]. However, understanding their roles in disease could lead to development of novel therapeutic targets [45]. For instance, *in vitro* evidence suggests that PAs have the capacity to facilitate protein aggregation [46–48]. However, the mechanism of action *in vivo* is not fully understood, specifically in pathological conditions where amyloidosis plays a critical role.

Here we hypothesized a relationship between PAs and Aβ accumulation, through the modulation of the ODC enzyme. Furthermore, we hypothesized that a dysregulation of the PA pathway contributes to Aβ oligomerization leading to neuronal dysfunction in DS-AD and AD. We utilized cell models of Aβ accumulation as well as brain tissue derived from normosomic and DS adults with AD and found that PA levels are increased in AD brain tissue, suggesting an interaction between PAs and Aβ_42_ aggregation. Additionally, we determined that ODC inhibition via the L-difluoromethylornithine (DFMO) inhibitor could reduce Aβ-mediated apoptosis as well as Aβ aggregation, thus potentially slowing AD pathology in a neuronal cell system.

## METHODS

### Reagents

Lipofectamine 2000, Lipofectamine 3000, Opti-MEM, Dulbecco’s Modified Eagle Medium (DMEM), were purchased from Invitrogen (Carlsbad, CA). Hank’s Balanced Salt Solution, Staurosporine, 2’,7’-Dichlorofluorescin diacetate (DCF), and Hoechst stain (St. Louis, MO). L-a-difluoromethylornithine was purchased from Cayman Chemical (Ann Arbor, MI). DsRed-Express2 plasmids were purchased from Clontech (Mountain View, CA). Primary antibodies for Amyloid-β Peptide antibody (MOAB-2) and Ornithine decarboxylase (ODC) antibody and cytosolic/mitochondrial tissue fractionation kit were purchased from Abcam (San Francisco, CA). Primary antibody for beta-actin was purchased from Cell Signaling Technologies (Danvars, MA). Cy3-conjugated secondary antibodies were purchased from Jackson Immunoresearch (Westgrove, PA). Normocin (Broad spectrum antimicrobial reagent) from InvivoGen (San Diego, CA). DL-α-Difluoromethylornithine (DFMO, CAS No. 96020-91-6), and PA standards were obtained from Cayman Chemical (Ann Arbor, MI). For more details on antibodies, see Table 1.

### Experimental Models

#### HT22 Cell Culture

Immortalized mouse hippocampal cells (HT22) were a gift from Dr. Russell Swerdlow’s research group (University of Kansas Medical School). They were plated in 35mm diameter 6-well culture plates in Dulbecco’s modified eagle’s medium (DMEM) containng 4.5g/L glucose supplemented with 10% fetal bovine serum (FBS), and penicillin (100 Units/mL)/streptomycin (100μg/mL). Cells were incubated overnight at 37°C in 10% CO_2_. The next day the HT22 cells were transfected and treated appropriately at 80-90% confluence.

#### H1b and Htk cell lines

The establishment of both H1b and Htk cell lines is reported elsewhere else [49], and were provided by Dr. Pablo Caviedes of the University of Chile. Cell lines were grown in 60 mm TPP tissue culture dishes and maintained in feeding medium, composed of DMEM/HAMF12 nutrient mixture (1:1) (Sigma Co., St. Louis, MO) supplemented with 6 g/L glucose, 1 g/L bicarbonate, 10% (v/v) adult bovine serum and 2.5% fetal bovine serum, and 100 µg/ml Normocin, 37C in 5% CO2. Passages were carried out by detaching the cells with Detachin^TM^ (Cell detachment solution, Genlantis, San Diego CA). The cultures were kept in incubators at 37°C, 100% humidity and an atmosphere of 5% CO2. Media was renewed twice a week.

#### Fibroblast Culture Maintenance and Treatments

Human fibroblasts were acquired from the Coriell Institute. Cells were grown primarily in DMEM media treated with Normocin (Invivogen) and appropriate concentrations of fetal bovine serum as recommended by their culture protocols (GM02036:GM02767). Cells in culture plates or wells were maintained in 5% CO_2_ for 37°C. During passaging at 90% confluence, cells were washed with phosphate-buffered saline (PBS), detached using Detachin (Genlantis), counted by hemacytometer, and resuspended in media. For treatments, DFMO was added to media prior to the cell resuspension at 0.1 mM.

#### Tissue Sources

The hippocampal brain tissue used was obtained from a collaboration with Dr. Ann-Charlotte Granholm and the MUSC Carroll Campbell Jr. Neuropathology laboratory. This South Carolina Brain Bank contains frozen and fixed tissues from >250 Alzheimer cases and Controls, and all cases have undergone neuropathological and clinical staging including the ABC assessment as described by Jack et al [50] and recently amended by DeTure and Dickson [51] and Aldecoa et al., [10]. Fifty milligram (mg) frozen tissue dissected from 3 neuropathologically-confirmed *postmortem* AD, DS-AD, and age-matched healthy control (HC) cases (Braak V-VI and CERAD C), and 3 age-matched controls which showed no AD pathology (Braak <II and CERAD 0-A) were collected (Supplementary Table 2). The tissue was then lysed and isolated into cytosolic and mitochondrial fractions according to the manufacturer’s protocol (Abcam, San Francisco, CA).

#### Plasmid Preparation

Empty pIRES2 DsRed-Express2 (DsRed2) bicistronic vectors containing Aβ_42_ sequence were originally transformed using 50 ng of plasmid in JM109 Escherichia coli and grown on LB agar plates containing 35 μg/mL kanamycin sulfate at 37°C overnight. Starter cultures were created using a colony containing transformed cells and grown in LB broth with 35 μg/mL kanamycin sulfate for 6-8 h at 37°C and diluted to 1:250 into overnight cultures. Plasmids were isolated and purified using a Maxi prep Kit according to the manufacturer’s instructions (Clontech, Mountain View, CA). Plasmid concentrations were quantified using three averaged recordings by a NanoDrop 2000 (Thermo Scientific). Bicistronic vectors that allow for co-expression of DsRed separate from the protein of interest were used in this experiment in place of protein fusion construction to prevent interference with the normal functioning and localization of the small proteins being examined. Because DsRed is expressed in the cell separately from Aβ_42_, there is no interference with normal function of Aβ_42_ or formation of unnatural aggregation following overexpression.

#### Transfection

HT22 cells were transfected with 5 μg, unless otherwise specified, of DsRed2 plasmids containing Aβ_42_ or empty IRES vector sequences using a standard Lipofectamine 2000 protocol as outlined by Bartley et al (2012). Both cell types were incubated in Opti-MEM with the DsRed2 bicistronic plasmid/Lipofectamine 2000 mixture for 4-6 h at 37°C in 10% CO_2_ for HT22. Transfection was terminated in both cell cultures after 4-6 h through replacement of the Opti-MEM with 1 mL culture media. Cells were incubated for 24 or 48 h at 37°C in 10% CO_2_ before proceeding with additional experimental analysis.

#### Fluorescence Staining and Imaging

Transfected and treated HT22 cells were washed once with PBS (pH 7.4) and fixed for 45 minutes in 4% paraformaldehyde at 25°C. Cells were washed twice with PBS then incubated in a 1:1000 dilution of Hoechst nuclear stain in PBS at 4°C overnight. The following day, cells were washed twice with PBS and placed in an anti-quench solution composed of 0.1% p-phenylenediamine in PBS for imaging. *All antibodies used in staining or Western Blot experiments are detailed in Supplemental Table 1*.

For immunohistochemistry experiments, following paraformaldehyde fixation, HT22 cells were washed twice with PBS then non-specific sites were blocked with 5% BSA in PBS containing 0.2% TritonX-100 for 1 h at 25°C. Blocking solution was then aspirated and cells were incubated overnight at 4°C in a primary antibody solution consisting of 2% BSA in PBS containing 0.2% TritonX-100 and primary antibody diluted according to manufacturer’s recommendations (Supplementary Table 1). Cells were then washed five times with PBS over the course of 1 h and then incubated in 1:1000 dilution of Hoechst nuclear stain and 1:250 dilution of secondary antibody in PBS at 25°C, protected from light for 1 h. Cells were then washed five times with PBS over the course of 1h before plating in an anti-quench solution composed of 0.1% p-phenylenediamine in PBS in for imaging.

Cells were imaged using a Zeiss Axiovert-200M epi-fluorescence microscope with five representative images taken of each well for assessment of apoptosis. All experimental conditions were performed in duplicate or triplicate wells in each experiment. Only cells fluorescing red through the expression of the DsRed-Express2 vector were counted and assessed for apoptosis through the visual evaluation of condensed or fragmented nuclei as indicated by Hoechst nuclear stain. Three wells with at least 25 transfected cells per wells were randomly selected and quantified for each experimental condition.

#### Reactive Oxygen Species (ROS) Detection

ROS detection was performed on live HT22 cells using DCF fluorescence 24 h post transfection. Anhydrous DCF was reconstituted in DMSO to yield 2mM stocks that were used for up to 2 months. Positive control groups were treated with H_2_O_2_ (125 μM) for 45-60 minutes at 37°C protected from light prior to loading with DCF. Then, all groups were loaded with 1 mL of 10 μM DCF dye in warmed PBS (pH 7.4) for 30 minutes at 37°C protected from light. Labeling solution was then aspirated and cells were stained with Hoechst stain (1μM) in warmed PBS for 15 minutes at 37°C protected from light. The cells were then mounted in anti-quench solution consisting of 0.1% p-phenylenediamine in PBS for imaging.

#### Thioflavin T (ThT)-Based Aggregation Kinetic Assays

Kinetic assays were conducted on an Infinite M200PRO plate reader (Tecan, Männedorf, Switzerland). Experiments were conducted in triplicate in a Costar black 96 well plate (Corning Inc., Kennebunk, ME). with a final volume of 200 μL per well. Every measurement was an average of 50 readings. The aggregation of Aβ_42_ was initiated by the addition of peptide from a stock solution (in DMSO, 0.5− 1 mM) to phosphate buffer to a final concentration of 10 μM. The concentration of ThT dye used in the experiment was 5 μM. The aggregation of Aβ_42_ was monitored by ThT fluorescence (λex = 445 nm and λem = 485 nm). The blank sample contained everything except Aβ_42_. The Kinetic assays in the presence of PAs were conducted under the matched conditions except that the PAs were added from a stock solution (10-50 mM in Milli Q water) to keep the final concentration of water less than 1.0% (v/v). The PAs were added to the wells with ThT dye and buffer and mixed gently before adding Aβ_42_. To keep the conditions identical, an equal amount of Milli Q water was added to the wells with Aβ_42_. Kinetic profiles were processed using Origin (version 9.1). The sample data were processed by subtracting the blank and renormalizing the fluorescence intensities of graphs and keeping the intensity of Aβ_42_ aggregation as one. The Kinetic curves of the Aβ_42_ aggregation were fit using the built-in sigmoidal fit. Each experiment was fit independently to extract the t_50_ (time required to reach 50% of the maximum fluorescence intensity).

#### HT22 Cell Lysis for MEK-LIF

HT22 cells were lysed either 24 or 48 h post transfection termination. First, cells were washed once with ice cold phosphate buffered saline solution (PBS, pH 7.4) before adding of Wahl lysis buffer (200 *µ*L) supplemented with leupeptin and aprotinin at volumes of 1 *µ*L per mL buffer, then incubated on ice for 10 minutes. Following incubation, a pipet tip was used to scrape the surface of each well before collection of lysis buffer. Each sample was centrifuged for 2 minutes at 9,600g and the resulting lysate was collected.

#### Micellar electrokinetic chromatography with laser induced fluorescence (MEKC-LIF): PA analysis

PAs were analyzed with a modified protocol from [52]. Briefly, samples were derivatized with 1.28 mM FITC/Acetone in 200 mM carbonate buffer (1:1) at pH 10 or derivatizing solution and stored at room temperature for 24 hours followed by analysis in the MEK-LIFD instrument.

#### MEKC-LIFD instrument

The MEKC-LIFD instrument was built in-house described elsewhere [53]. In brief, a Glassman High Power Supply model PS-MJ30P04400X88 (Glassman High Voltage Inc. Whitehouse Station, NJ, USA), a collinear fluorescence detector described elsewhere [54] equipped with a Hamamatsu Photo Multiplier Tube Model H-9306 (Hamamatsu Co, Middlesex, NJ, USA) and a Cobolt state, 488 nm, continuous wave laser model 0488-06-01-0060-100 (Cobolt MLD, Solna, Sweden). The signals were collected by a National Instruments data acquisition card, model PCI-6221 (National Instruments Co. Austin, TX, USA) in a computer programmed with custom-made software both for acquisition and processing.

#### Sample treatment

*Cell lysates:* After quantifying protein concentration via BCA protein assay, the cell lysates were deproteinized by mixing lysates with acetonitrile in a ratio of 1:1, vortex and centrifuged at 14,000 g at 4 °C for 15 minutes. Supernatants were mixed with derivatizing solution (1:1). After 24 h in darkness, the samples were ready for MEKC-LIF analysis. To identify PAs, samples were spiked with a solution of 2 μM of PA standards and derivatized as described above. Then the electropherograms were compared by overlapping to detect the enhanced peak. *Brain tissue*: tissue was homogenized in 20 mM borate buffer (pH 9.3) in a ratio of 2 mg of tissue per 100 uL of buffer. The homogenates were mixed with an equal volume of chloroform and vortexed (20 min) followed by centrifugation at 15,000 g for 20 min, the supernatants were transferred to Eppendorf tubes and deproteinized with acetonitrile at 1:1 ratio, followed by derivatization as described above. Samples were centrifuged at 14,000 g for 15 minutes and stored in dark at room temperature for 24 hours, after which they were ran in the MEKC-LIFD for quantification of PAs.

#### Capillary electrophoresis

The standard solutions and the samples were hydrodynamically injected by applying a negative pressure of −10 psi for 1 s at the cathodic end of a fused silica capillary model TPS 025375, polyimide coated, 350 μm external diameter, 25 μm inside diameter and 60 cm long (Polymicro Technologies Inc., Phoenix, AZ, USA) while the anodic end of the capillary was immersed into the tested solution. The running buffer was 40 mM Sodium Borate and 20 mM SDS at pH 9.0 [55]. After injection the anodic end of the capillary was immersed in a buffer reservoir equipped with a Platinum-Iridium electrode while the cathodic end remained in the buffer reservoir of the cathode. A positive, 27 kV was applied at the anode for the electrophoretic separation and the data collected for 45 minutes.

#### Sample data analysis

The signal was collected at 40 points per second using a homemade software in MATLAB® and data output in the form of voltage versus time was presented as electropherograms. The data were analyzed by means of a PicoAnalytical software Inc. (Miami FL), that is capable of filtering with a moving average digital filter and has both horizontal and vertical displacement capabilities for automatic electropherograms alignment. The software deconvoluted the peaks and calculated the best fitting Gaussian curve for each peak. The same software also calculated the average of unlimited number of electropherograms.

#### Cell Lysis for Western Blotting and A***β***_42_ ELISA

HT22 and human fibroblast cells were lysated at either 24 or 48 h post transfection termination. First, cells were trypsinized before being collected and centrifuged for 5 minutes at 1100 g. The pellets were then washed with ice cold Dulbecco’s phosphate buffered saline solution (DPBS) before centrifuged again for 5 minutes at 3,500 g. The resulting pellets were resuspended in 1X RIPA buffer (150 mM NaCl, 1% NP-40, 0.5% sodium deoxycholate, 0.1% SDS, 50 mM Tris, pH 8.0) supplemented with leupeptin and PMSF and incubated on ice for 30 minutes. The samples were then sonicated in a water bath for 1 minute before being centrifuged at 14,000 g for 15 minutes at 4°C. The subsequent supernatants were collected for analysis.

#### Western Blotting

Western blots were utilized to analyze HT22 and human fibroblast cell lysates (See Cell Lysis for Western Blotting and Aβ_42_ ELISA). Protein samples (10-15 µg/lane) were resolved by 8-16% gradient Novex Tris-glycine gels (Thermo Fisher, Carlsbad, CA) and proteins were then transferred to PVDF membranes. Non-specific binding sites were blocked using 1% BSA in PBS (pH 7.4) containing 0.1% Tween-20 (PBS-T) for 1 h at 25°C. *All antibodies used for Western blotting are detailed in Supplemental Table 1.* The blocking buffer was drained, and the membrane was allowed to incubate in primary antibody diluted in blocking buffer according to manufacturer’s recommendations (Supplementary Table 1) overnight at 4 °C. The membrane was washed three times for 15 minutes in PBS-T and was then incubated with the secondary antibody for 1.5 h at 25 °C. The secondary was then removed, and the membrane was washed again in PBS-T, three times for 15 minutes. Immunoreactive proteins were detected using enhanced chemiluminescence (ECL) reagents (GE Healthcare; Pittsburgh, PA) and films were imaged using a ChemiDoc (BioRad, Hercules, CA). Re-probing of blots was performed by stripping in 0.1 M Tris-HCl (pH 8.0), 2% SDS, and 100 mM β-mercaptoethanol for 30 minutes at 52 °C. The blots were rinsed twice in PBS-T and processed as above with a different primary antibody.

#### DFMO Treatment

Six hours following plating, HT22 cells were pre-treated with difluoromethylornithine (DFMO) (5 mM; stock made in culture media) before transfection. Cells were transfected according to the protocol listed. Following the removal of the transfection medium, DFMO (5 mM) was administered. All cells were left in treatment media for 24 or 48 h before proceeding. HtK and H1B cells were cultured in growth media containing DFMO 5 mM throughout three consecutive passages, after which these cells were analyzed for Aβ_42_ monomers (ELISA), proteostat and quantification of PAs.). H1b and Htk cells (200,000 cells/mL) were plated in 35 mM sterilized petri dishes and incubated at 37 °C and 5% CO_2_ (g) and allowed to adhere to the plate for 24 h in media (DMEM, 10% FBS, 1% pen/strep). The cells were detached from culture flasks using detachin (manufacturing specs) and collected before centrifugation for five minutes at a rate of 1,500 g. These cells (HtK and H1b, 1×10^6^ cell/mL) were used to seed plates in the absence or presence of DFMO (5 mM final concentration in cell media). The cells were incubated for 48 h. Subsequently, the cells were detached and passaged again and treated them with DFMO under the matched conditions as described above. The cells were further incubated for 72 h and then detached from plates using detaching and collected before centrifuging for 5 minutes at a rate of 1,500 g. All subsequent centrifugation steps were performed in the same manner.

#### Aβ_42_ Monomer ELISA

Monomeric Aβ_42_ was detected in both HT22, H1b, and Htk cell lysates utilizing an Aβ_42_ Mouse enzyme-linked immunosorbent assay kit (ELISA) (Thermo Fisher, cat #KMB3441, Carlsbad, CA). Briefly, Aβ_42_standard was reconstituted in sodium bicarbonate (55 mM, pH 9.0) then utilized to generate a standard curve ranging from 3.12 – 200 pg/mL in a 96 well pre-coated clear bottom plate. Next, 100 µL of each sample was plated in duplicate and allowed to incubate at room temperature for 2 h. Following incubation, wells were washed four times with 1X wash buffer before adding 100 µL of detection antibody, which was incubated at room temperature for 1 h. The wells were then washed four times with 1X wash buffer before adding 100 µL Streptavidin-HRP and incubating at room temperature for 1 h. Then wells were then washed four times with 1X wash buffer before adding 100 µL chromogen to each well and incubating protected from light at room temperature for 30 minutes. Finally, 100 µL stop solution was then added to each well before reading with a colorimetric microplate reader at 450 nm.

#### Proteostat Dye Assay

H1b and Htk cell lines were analyzed for intracellular protein aggregation using the ProteoStat Protein Aggregation Assay Kit (Enzo Life Sciences, Farmingdale, NY). Once the cells were treated with DFMO (as described above), they were detached form plates using detachin and collected before centrifugation for five minutes at a rate of 1,500 g. The pellet was then homogenized using 1 mL PBS (pH 7.4) before centrifuging again. The resulting pellet was fixed using 4% paraformaldehyde for 30 minutes on ice. The samples were then centrifuged again, and the resulting pellet was then homogenized using 1mL PBS (pH 7.4) before centrifuging again. Subsequently, 500 μL 0.1% Triton was added to each sample and incubated for 20 minutes on ice. The samples were then centrifuged again, and the resulting pellet was then homogenized using 1 mL PBS (pH 7.4) before centrifuging again. A total of 375 μL of proteostat dye (0.001% proteostat dye in 1X PBS) was then added to each sample and incubated protected from light for 20 minutes at room temperature. The samples were then centrifuged, then homogenized with 400 μL PBS. The cells were then transferred to a black 96 well plate (Corning Inc., Corning, NY) (100 µL/each well and four wells per condition) and the fluorescence was measured using a 96-well plate reader (λ_ex_= 550 nm, λ_em_= 600 nm). The experiments were conducted at least three times and four technical replicates were conducted for each experiment. The reported error bars for ProteoStat fluorescence intensity are the s.d.’s for three sets of experiments.

#### Cell Viability Assay

Fibroblasts and mouse cells were treated four days or 24 hours prior to cell viability analysis. Cells were measured for viability primarily using MTT assays. Initial cell density was determined using Trypan Blue with an automatic cell counter and treated with DFMO when specified. Additional wells were treated with dimethyl sulfoxide (DMSO) as a control. On the day of analysis, the media was exchanged for OPTIMEM (Gibco, 31985062) containing MTT to limit further cell growth while providing sufficient conditions for a three-hour incubation. After incubation, the crystals formed were dissolved in DMSO and read for absorbance at 570 nm.

#### Immunofluorescence of Cell Culture and Tissue

Fibroblasts were grown in 4-well chambers (Cellvis, C4-1.5H-N) pre-treated with 0.1mg/mL. After fixation by 4% paraformaldehyde/4% glucose fixative, cells were stained with the indicated primary antibodies targeting selected proteins overnight followed by fluorophore-conjugated secondaries. Paraffin-embedded human hippocampal tissue were graciously provided by Dr. Lotta Granholm’s lab from the University of Colorado Anschutz. Tissues were deparaffinized, quenched for autofluorescence (Biotium, #23007) and treated with the indicated primary antibodies for an overnight incubation. Secondaries were introduced during the second day followed by ProLong Gold mounting overnight at −20°C. Cells and tissues were imaged under an Olympus confocal microscope.

#### Experimental Design and Statistical Analysis

All experimental conditions were performed in duplicate and statistical analysis was only performed on experiments quantified a minimum of three times. Data represents the means +/-SEM for the total number (n) of experiments carried out. For statistical comparisons, an analysis of variance (ANOVA) was carried out followed by a post hoc test Tukey’s test for individual comparisons, using GraphPad Prism 10 software. A p-value of <0.05 was considered statistically significant.

## RESULTS

### PA levels in AD tissues

We examined PA levels in *postmortem* hippocampal tissue from AD patients. Using MEKC-LIF, *postmortem* hippocampal samples from patients with a confirmed neuropathological diagnosis of AD or DS-AD as well as age-matched controls were analyzed for levels of putrescine, spermidine, and spermine (Figure 1A-C). Levels of spermidine [t(4)=4.989, p=0.0075] and spermine [t(4)=3.166, p=0.034] were significantly elevated in samples from individuals with AD in comparison to age-matched control samples (n= 3 per group). These findings confirm previous and current (see below) findings in AD mouse and cell models.

**Figure 1:**
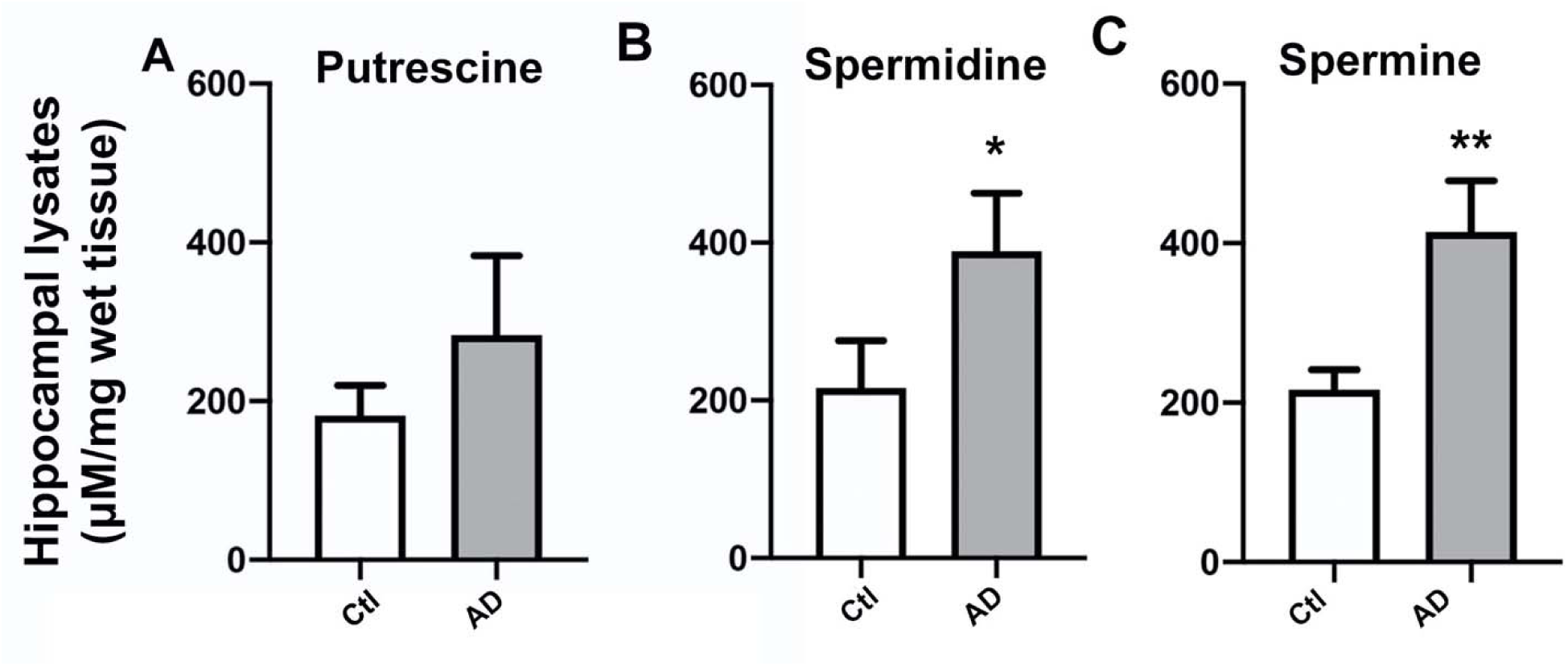
PA levels were increased in AD hippocampal tissue samples. MEKC-LIF was used to measure concentrations of putrescine **(A),** spermidine **(B),** and spermine **(C)** in lysates of postmortem human hippocampal tissue of sporadic AD and age matched controls. Results shown as mean SEM, n=3. ** indicates p<0.01 compared to control group; * indicates p<0.05 compared to the control group as determined using t-test with a post hoc Tukey’s test.

### Putrescine accelerated Aβ_42_ in-vitro

To confirm whether the PA putrescine could accelerate aggregation of Aβ*_42_*, we first utilized a cell free-Tht fluorescence aggregation assay (Figure 2A-B). Figure 2A depicts amyloid formation kinetics in the ThT fluorescence assay over time for 10 μM Aβ*_42_* peptide in the presence (black) or absence (red) of putrescine (100 μM). In the presence of putrescine, aggregation was faster. Aβ_42_ aggregation relative intensity is displayed in black in the presence of putrescine at 50:1 molar ratio and in red without putrescine. Comparing the time to reach 50% fluorescence (t_50_) for Aβ_42_ showed a dose-dependent increase in aggregation speed as a function of putrescine concentration. Therefore, in a cell-free environment, PAs increased the speed of Aβ*_42_* aggregation.

**Figure 2:**
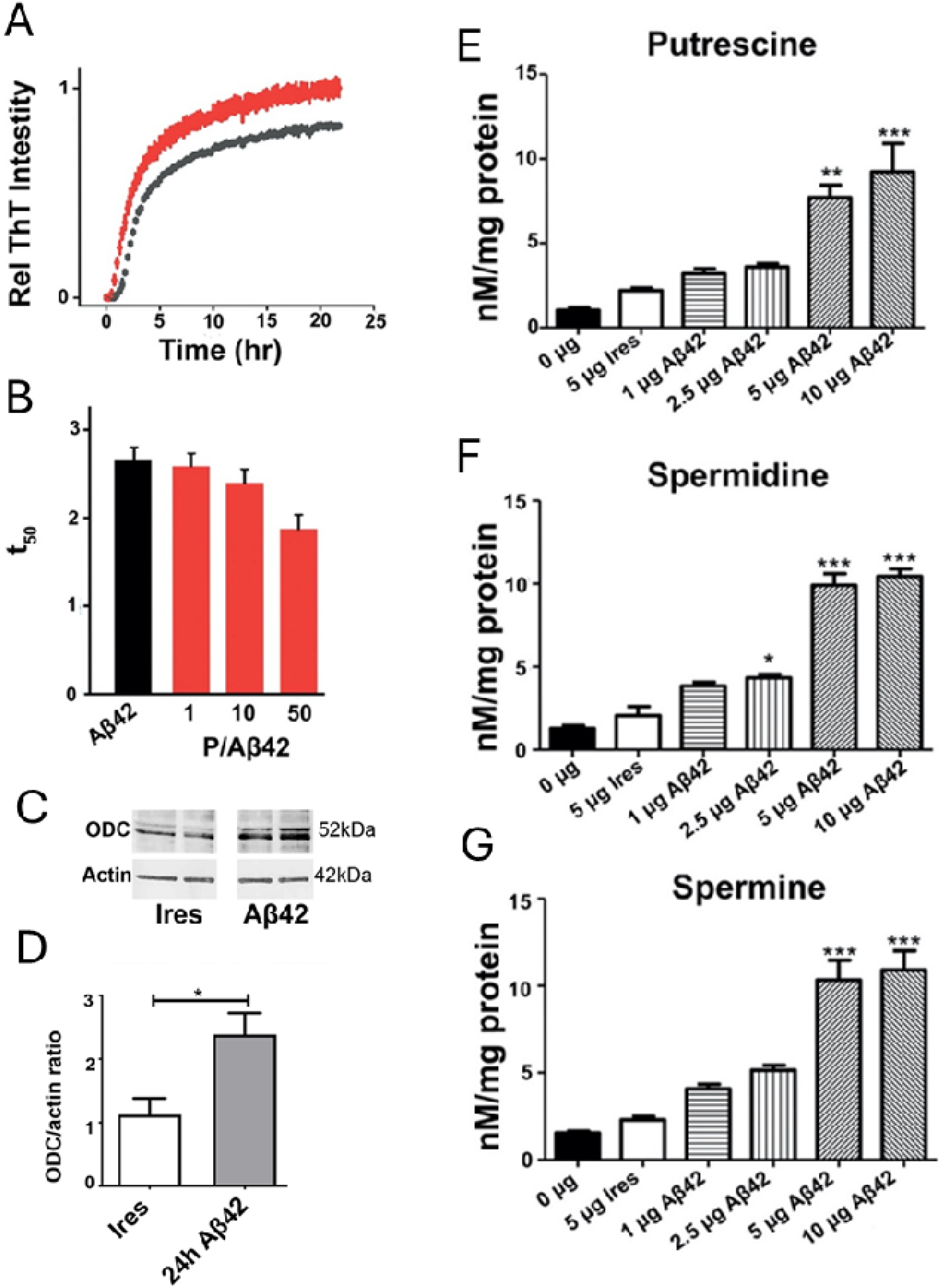
PA levels increased with increasing concentration of Aβ_42_. **A & B)** ThT fluorescence assays. Amyloid formation kinetics of 10 μM Aβ1-42 peptide in the presence or absence of putrescine (100 μM). Average of triplicate wells with standard deviation plotted. **A)** ThT fluorescence assay of Aβ_42_ over time. Aβ_42_ aggregation relative intensity is displayed in black in the presence of putrescine at 50:1 molar ratio. **B)** Tht fluorescence assay time to reach 50% fluorescence (t50) for Aβ_42_ is indicated in black versus t50 for various molar ratios of putrescine as follows; 1:1, 10:1, 50:1 displayed in red. **C)** Western blot analysis of HT22 lysates acquired 24 hours post-transfection after transfection with empty pIRES DsRed-Express2 (Ires) bicistronic vector or vector co-expressing DsRed2 with Aβ_42_ using antibodies specific for ODC (top panel) and beta-actin (bottom panel). **D)** Densitometry quantification of (A) and (B). **E-F)** MEKC-LIF was used to measure concentrations of putrescine **(E)**, spermidine **(F)**, and spermine **(G)** in HT22 cell lysate that was acquired 48 hours after transfection with no plasmid (0 μg), empty pIRES DsRed-Express2 (Ires) bicistronic vector (5 μg) or vector co-expressing DsRed2 with Aβ_42_at increasing concentrations (1, 2.5, 5, or 10 μg). Results shown as mean SEM, n=3. ** indicates p<0.01 compared to IRES control, *** indicates p<0.001 compared to IRES control as determined using one-way ANOVA with a post hoc Tukey’s test.

### Aβ_42_ induced ODC expression in HT22 cells

One possible mechanism for increased PA levels in the presence of abundant Aβ*_42_* could be that expressed Aβ*_42_* promotes the activity of the rate-limiting PA-producing enzyme ODC. To test this possibility, lysates of 24-hour IRES and Aβ*_42_*transfected HT22 cells were analyzed using Western blotting with an antibody specific to ODC. As shown in Figure 2C, the bands detecting ODC at 52 kDA were noticeably darker in the Aβ*_42_* sample lysates in comparison to the IRES samples. After normalizing to beta-actin, one-way ANOVA revealed a significant increase in the ratio of ODC abundance 24 hours post transfection with Aβ*_42_* in comparison to those with IRES as well as 48 hours post transfection [F(2,13)=9.207, p=0.0032] (Figure 2D). We therefore concluded that the Aβ*_42_* transfected cells had increased levels of ODC post transfection as compared to control HT22 cells.

### Aβ_42_ aggregation and polyamine levels

We then evaluated whether Aβ*_42_* aggregation could cause changes in PA levels in hippocampal neurons. HT22 cells were transfected with either empty IRES vector or increasing concentrations of Aβ*_42_*for 48 hours. Subsequent analysis of the 48-hour cell lysates using MEKC-LIF yielded a dose dependent increase in which higher transfection concentrations of Aβ*_42_* were associated with similarly increased levels of the PAs, putrescine, spermidine and spermine, in HT22 cells (Figure 2E-G). Separate one-way Analysis of Variance (ANOVA) were conducted for putrescine, spermidine and spermine followed by post hoc analysis to isolate simple effects. As can be seen in Figure 2C there was a significant main effect of transfection dose on putrescine levels, with 5 μg and 10 μg of Aβ*_42_*producing significantly higher levels of putrescine compared to 0 μg of Aβ*_42_*, 5 μg of IRES, 1 μg of Aβ*_42_*, and 2.5 μg of Aβ*_42_* [F(5,12)=17.93, p<0.0001]. Similarly, there was a significant main effect of transfection dose on spermidine levels, where transfection with every dose of Aβ*_42_* elicited significantly higher spermidine levels compared to 0 μg of Aβ*_42_* [F(5,12)=82.35, p<0.0001]. Additionally, transfection with 2.5, 5 and 10 μg of Aβ*_42_*produced significantly higher levels of spermidine compared to 5 μg of IRES (p<0.05). A significant dose response of Aβ*_42_* transfection was also shown with spermine [F(5,12)=35.55, p<0.0001], where 5 and 10 μg of Aβ*_42_* producing significantly higher levels of putrescine compared to 0 μg of Aβ*_42_*, 5 μg of IRES, 1 μg of Aβ*_42_*, and 2.5 μg of Aβ*_42_*. Notably, transfection with 5 and 10 μg of Aβ*_42_* caused a significant increase in the concentration of all three PAs as compared to IRES, with putrescine showing the largest dose-dependent relationship between Aβ*_42_*and upregulation. These data indicated that while PAs are shown to influence Aβ*_42_* aggregation, Aβ*_42_* can also in turn affect PA production.

### DFMO decreased PA levels in response to Aβ_42_

Difluoromethylornithine (DFMO) is an irreversible inhibitor of ODC. As PAs affected Aβ*_42_* aggregation speed, we hypothesized that inhibiting ODC could affect Aβ*_42_* aggregation and subsequent effects. First, we investigated whether DFMO could reduce the increase in PA induced by Aβ*_42_*. HT22 cells were pre-treated 6 hours after plating with DFMO (5 mM) in growth media, where they were incubated overnight before transfection the following day. When terminating transfection, the cells were treated again with DFMO (5 mM) in growth media followed by incubation. Using MEKC-LIF, lysates collected 48 hours post transfection were analyzed for PA levels (Figure 3). Separate one-way ANOVAs were conducted for putrescine, spermidine and spermine followed by post hoc analysis to isolate simple effects. As can be seen in Figure 3, there was a significant main effect of Aβ*_42_* transfection on putrescine levels [F(2,7)=20.16, p=0.0012], spermidine levels [F(2,7)=47.97, p<0.0001] and spermine levels [F(2,7)=41.56, p<0.0001]. Specifically, putrescine, spermidine and spermine levels were significantly higher after Aβ*_42_* transfection compared to IRES, and the increase in PAs was attenuated by DFMO. Therefore, the inhibition of ODC by DFMO reduced PA production due to Aβ*_42_*.

**Figure 3:**
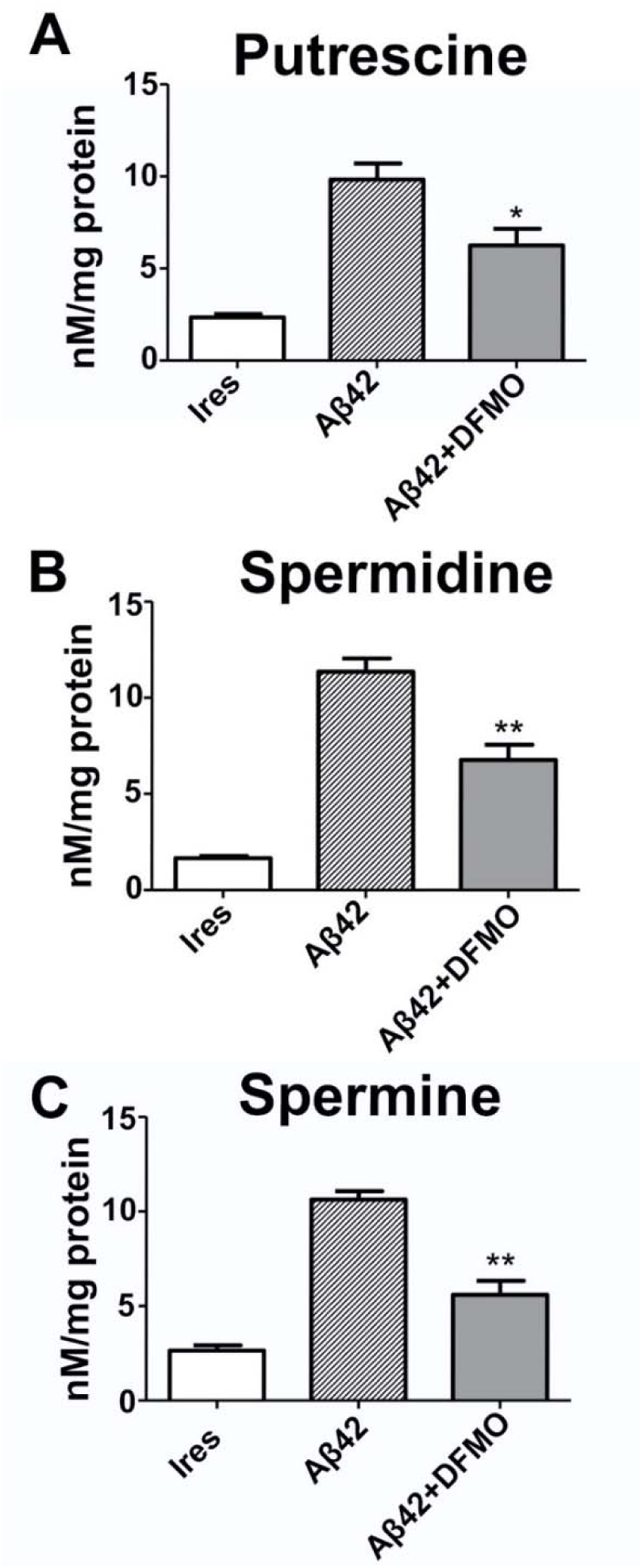
ODC-inhibitor DFMO decreased Aβ_42_-induced PA levels. MECK-LIF was used to measure concentrations of putrescine **(A)**, spermidine **(B)**, and spermine **(C)** in HT22 cell lysate that was acquired 48 hours after transfection with either empty pIRES DsRed-Express2 (Ires) bicistronic vector or vector co-expressing DsRed2 with Aβ_42_ at in the presence or absence of difluoromethylornithine (DFMO; 5mM). Results shown as mean SEM, n=3 for Ires and Aβ_42_, n=4 for Aβ_42_ + DFMO. * indicates p<0.05 Aβ_42_+DFMO compared to Aβ_42_ alone, ** indicates p<0.001 Aβ_42_+DFMO compared to Aβ_42_alone as determined using one-way ANOVA with a post hoc Tukey’s test.

### DFMO promoted Aβ_42_ monomer presence and attenuated cellular stress in HT22 cells

Next, we analyzed the effects of ODC inhibition by DFMO and thus reduced levels of PA on Aβ*_42_*aggregation in HT22 cells. Imaging of the cells following transfection and treatment protocol revealed that DFMO decreased the relative amount of Aβ_42_ aggregation in Aβ_42_ transfected cells, as evidenced by noticeably less intense aggregate staining in the Aβ_42_ + DFMO images in comparison to those without DFMO (Figure 4A). Further, using an ELISA assay, we measured the levels of Aβ_42_ monomers 48 hours post transfection. As can be seen in Figure 4B, a one-way ANOVA revealed a significant increase in monomers in cells treated with DFMO as compared to Aβ*_42_* and IRES, suggesting a reduction in aggregated peptides in the treated group [F(2,9)=5.299, p=0.0301].

**Figure 4:**
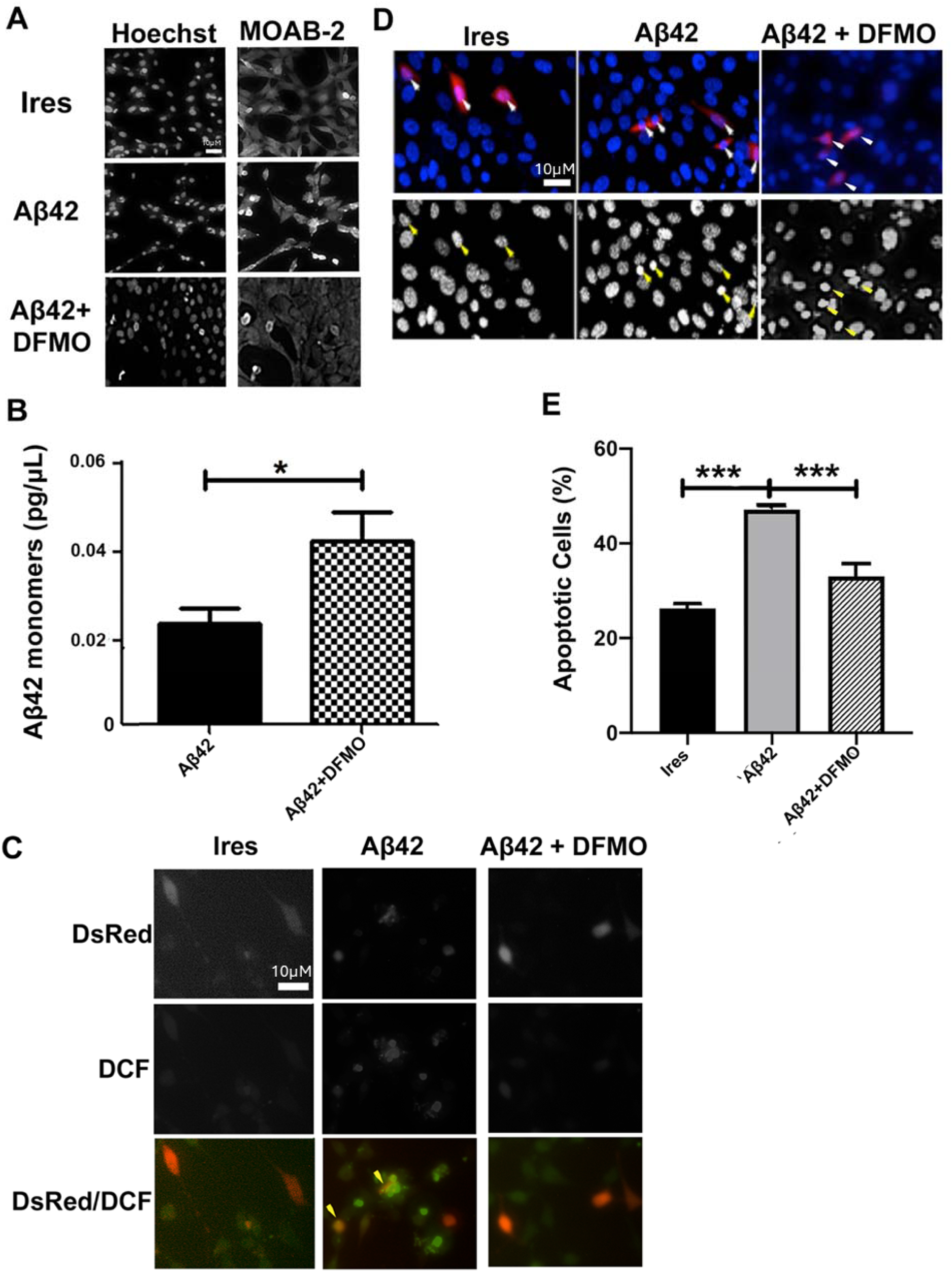
DFMO decreased Aβ_42_ aggregation, ROS production, and apoptosis *in vitro*. **A)** Representative HT22 cells imaged 48 hours after transfection with empty pIRES DsRed-Express2 (Ires) bicistronic vector or vector co-expressing DsRed2 with Aβ_42_ in the presence or absence of difluoromethylornithine (DFMO; 5mM). On the left, decolorized nuclei are shown stained with Hoechst dye. On the right, decolorized Aβ_42_ aggregates are shown. Scale bar = 10 microns. **B**) ELISA quantification of Aβ_42_ monomers in lysates from HT22 cells 48 hours after transfection with bicistronic vector co-expressing DsRed2 with Aβ_42_ in the presence or absence of DFMO (5 mM). Results shown as mean SEM, n=4. * indicates p<0.05 Aβ_42_+DFMO compared to Aβ_42_ alone**. C)** Representative images of DCF staining assay in HT22 cells 48 hours after transfection as described in **(A).** Successfully transfected cells are indicated by DsRed2 fluorescence in the top panel, oxidative species are indicated by DCF staining in the middle panel. Overlay of the two signals are shown in the bottom panel. Arrows indicate transfected cells with positive ROS staining. Scale bar = 10 microns. **D)** Representative images of HT22 cells imaged 48 hours after transfection with IRES bicistronic vector co-expressing DsRed2 with Aβ_42_. in the presence or absence of DFMO (5 mM). Arrows indicate transfected cells through DsRed2 fluorescence (red), nuclei are stained with Hoechst dye (blue). Bottom panels are shown in black and white to emphasize nuclear morphology. Scale bar = 10 microns. **E)** Quantification of apoptosis observed in HT22 cells 48 hours after transfection with empty IRES bicistronic vector or vector co-expressing DsRed2 with Aβ_42_ in the presence or absence of DFMO (5mM). Only DsRed2-positive cells were counted and marked as apoptosis based on the appearance of condensed or fragmented nuclei as indicated by Hoechst nuclear stain. Data are expressed as the mean SEM, n=4, independent experiments performed in triplicate. *** indicates p<.001 Aβ_42_ compared to IRES control; Aβ_42_ + DFMO compared to Aβ_42_ alone as determined using a one-way ANOVA with a post hoc Tukey’s test.

Aβ expression and aggregation is known to cause increased ROS levels. With a decreased level of Aβ42 aggregation with ODC inhibition, it is also possible that ROS levels would be similarly reduced. We next measured the effects of DFMO on Aβ-induced ROS using DCF staining. ROS was noticeably reduced in cells treated with DFMO, as displayed through the reduction in DCF signal (Figure 4C).

Since reduction in the PA synthesis reduced both Aβ aggregation and associated ROS, we sought to determine if such inhibition would reduce apoptosis in Aβ_42_-transfected cells. Upon quantifying the percentage of apoptotic transfected cells 48 hours post transfection marked by the DsRed signal and nuclear condensation and fragmentation, there was a significant increase in apoptosis in cells transfected with Aβ*_42_* in comparison to IRES [F(2,9)=43.83, p<0.0001]. Interestingly, treatment with DFMO significantly reduced the number of apoptotic cells following Aβ*_42_* transfection (Figure 4E). Taken together, these data support that application of DFMO in physiological doses partially protected HT22 cells from Aβ*_42_* aggregation, and in turn, ROS production and cell death.

### DFMO decreased Aβ_42_ aggregation in the Htk trisomy cell line

Htk cells harbor a triplication of APP, compared to the disomic cells obtained from their wild type littermates’ H1b cells. Aβ_42_ aggregates are markedly increased in Htk cells in comparison to H1b cells as shown through MOAB-2 staining (Figure 5A). To investigate if the inhibition of ODC could decrease aggregation in the Htk cell line, we treated the cells with 5 mM DFMO and measured aggregation with the Proteostat dye, which is a generalized marker for protein aggregation. As can be seen in Figure 5B, there was a significant decrease in aggregation in Htk cells treated with DFMO compared to those without treatment [t(4)=5.107, p=0.0069]. Two way analysis of the cell lines (Htk, H1b) and treatment (vehicle, DFMO) failed to detect a significant interaction, however there was a significant main effect of cell line with Htk having higher Aβ_42_monomers compared to DFMO treatment [F(1,12)=6.566, p=0.0249]. Next, the PAs, putrescine, spermidine, and spermine were measured in the H1B and Htk DS cell lines and DFMO was applied to the cultures to assess whether PAs in the DS cell cultures were altered (Figure 5C-E). Separate one-way ANOVAs were conducted for putrescine, spermidine, and spermine followed by posthoc analyses to isolate effects. Putrescine [F(2,21)=18.71, p<0.0001], spermidine [F(2,21)=5.814, p=0.0098] and spermine [F(2,21)=9.248, p=0.0013] were elevated in the Htk cell line compared to the H1b control cell line. Additionally, DFMO significantly reduced the levels of putrescine, spermidine and spermine in the Htk DS cell line. These data suggest that modulating PA synthesis by inhibiting ODC poses a viable approach for reducing Aβ_42_ aggregation, both in AD in the general population and in those with DS-AD. Interestingly, the drug affected the aggregation process both in cells that had an artificially added or transfected with Aβ_42_and in naturally APP over-expressing cells (Htk cells), providing increased scientific rigor for its validity as an amyloid-reducing drug.

**Figure 5:**
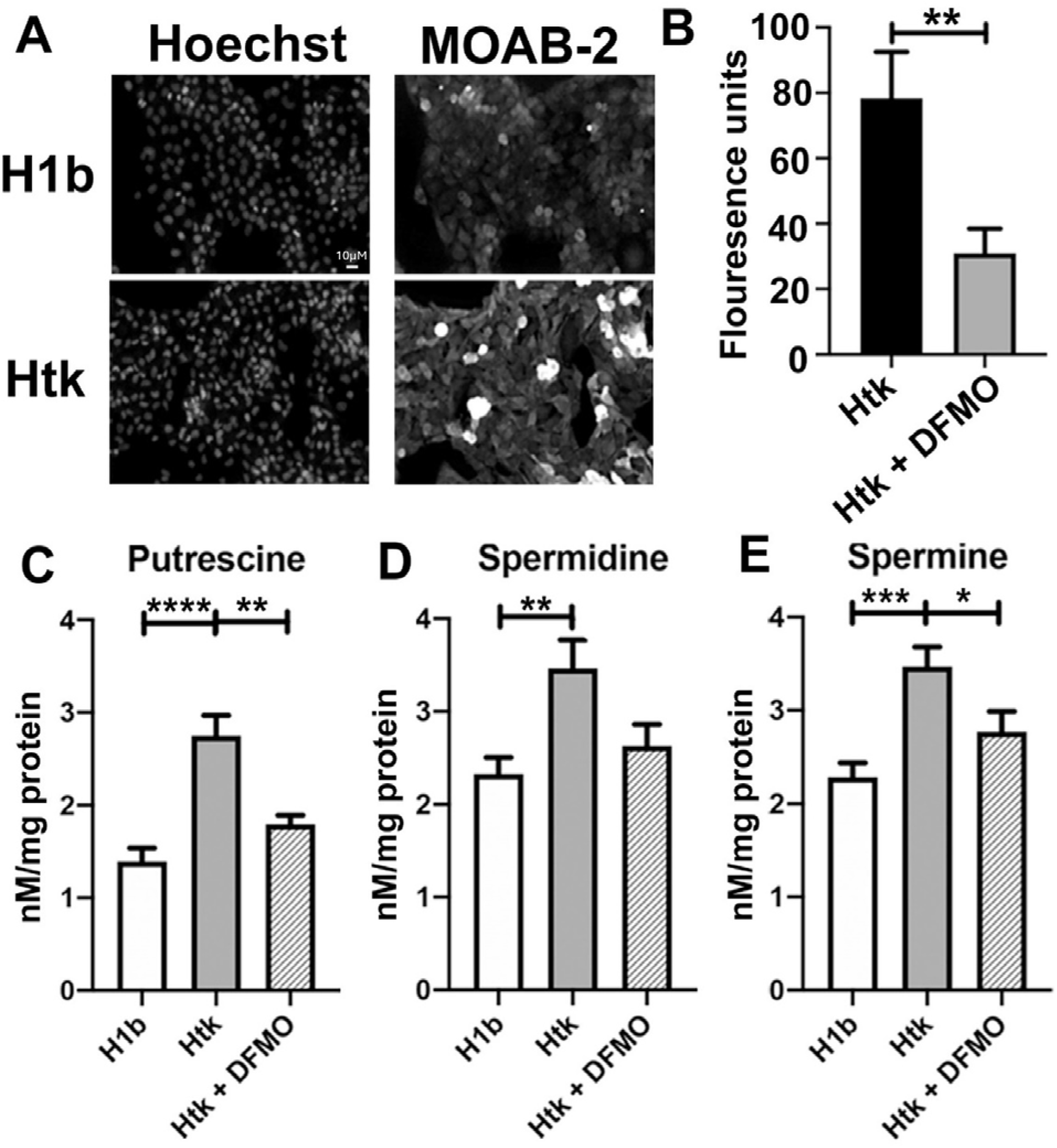
DFMO decreased Aβ_42_ aggregation in Htk cell line. **A)** Representative images of Aβ_42_aggregation hippocampal mouse cell line (H1b), hippocampal mouse trisomic Down Syndrome cell line (Htk). On the left, decolorized nuclei are shown stained with Hoechst dye (DAPI). On the right, decolorized Aβ_42_ aggregates are shown (MOAB-2). Scale bar = 10 microns. **B)** Proteostat analysis of aggregation in Htk cell lines in the presence or absence of difluoromethylornithine (DFMO; 5 mM). Data are expressed as the mean SEM, n=2. **C-E**) MECK-LIF was used to measure concentrations of putrescine **(C),** spermidine **(D),** and spermine **(E)** in lysates collected from H1b and Htk cells in the presence or absence of difluoromethylornithine (DFMO; 5mM). Results shown as mean SEM, n=8. **** indicates p<0.0001,*** indicated p<0.0001, ** indicates p<0.001, * indicates p<0.05 as determined using one-way ANOVA with a post hoc Tukey’s test.

### Effects of DFMO Treatment in normosomic and trisomic Human Fibroblasts

.Densitometry revealed significant changes in relative protein concentrations both between the normosomic and trisomic fibroblasts as well as in cell cultures treated with 5.0 mM DFMO for 48 hours. Arginase 1 did not show significant changes between karyotype or treatment (Figure 6A), but ODC levels were reduced in the T21 culture compared to untreated D21 cells (Figure 6B). For PA biosynthesis enzymes, T21 had increased levels of spermine (Figure 6C) and spermidine synthases (Figure 6D). With the addition of DFMO, the concentrations of both enzymes reacted similarly, with DFMO increasing spermine synthase in D21 (not significantly in the case of spermidine synthase) and a decrease in the levels of both enzymes in the DFMO treated T21 cells. Spermine/spermidine acetyltransferase 1 (SAT1) was not significantly affected by the addition of DFMO, nor was it altered significantly in T21 fibroblasts (Figure 6E). BACE1 levels were significantly higher in T21 fibroblasts compared to D21 control cells, and the addition of DFMO gave rise to a significant reduction of BACE levels in T21 cells (Figure 6F).

**Figure 6:**
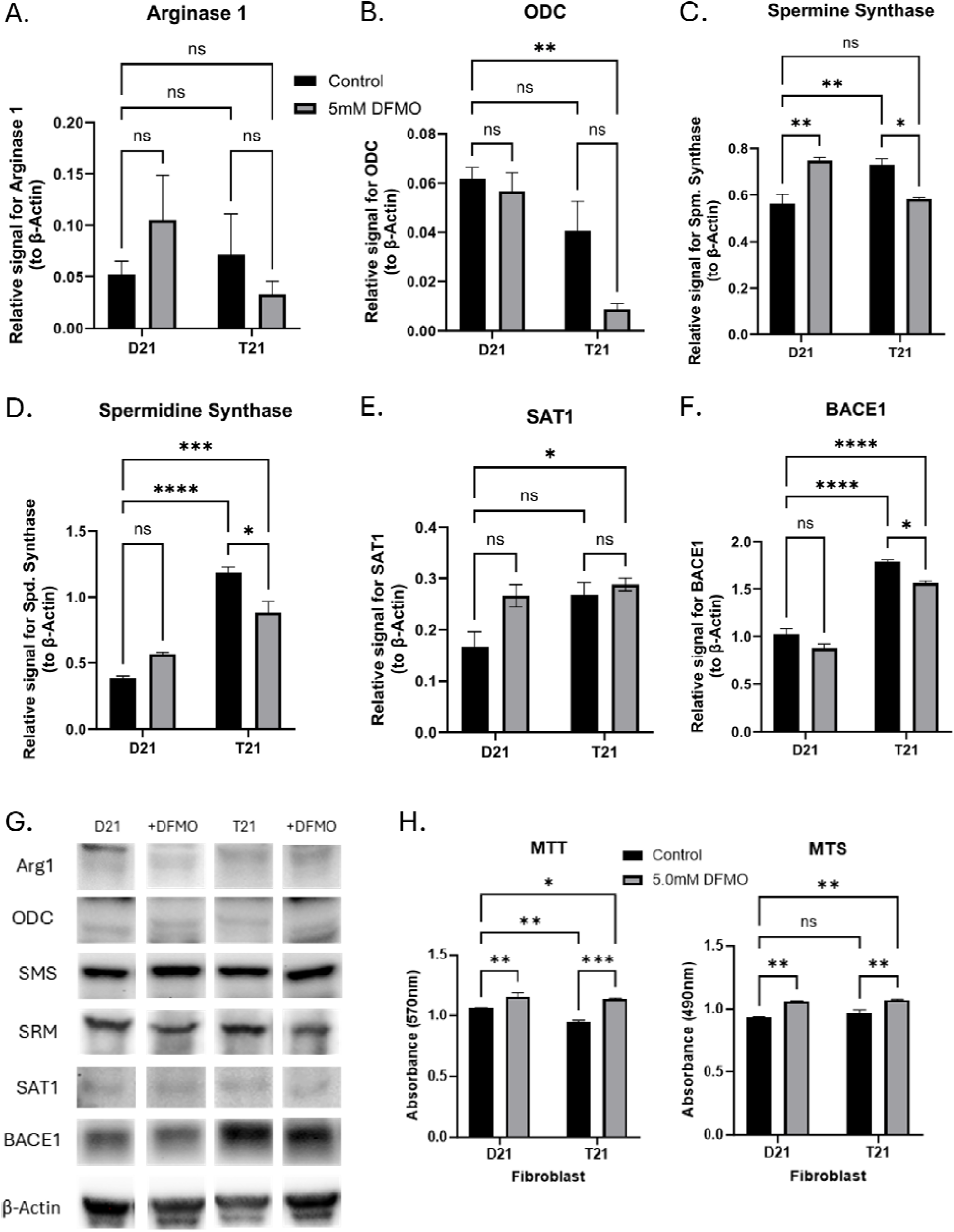
Effect of DFMO Treatment on PA Metabolic Enzymes and Cell Viability in Normosomic and Trisomic Human Fibroblast. Western blot densitometry of normosomic (D21) and trisomic (T21) human fibroblast cell culture treated with 5.0 mM DFMO for 48 hours before lysis (n=3), targeting: A) arginase 1, B) ODC, C) spermine synthase, D) spermidine synthase, E) SAT1, F) BACE1 with B-Actin control. G) Representative images of the indicated proteins. H) MTT and MTS analysis of fibroblast cells treated with 5mM DFMO for 48 hours. Results shown as mean SEM, n=3. **** indicates p < 0.0001, *** indicated p < 0.001, ** indicates p < 0.01, * indicates p < 0.05 as determined using two-way ANOVA with a post hoc Tukey’s test.

We employed MTT and MTS assays to observe the effects of DFMO on cell viability (Figure 6H). T21 cells were significantly reduced in absorbance of MTT crystals but not MTS. When treated with 5.0 mM DFMO 48 hours prior to experimentation, there was an increase in cell viability in both T21 and D21 cell lines.

### ODC Expression and correlation with APP

Human fibroblasts were immunostained for amyloid precursor protein (APP) and ODC and densitometry was performed for each protein with confocal microscopy (Figure 7). Only in D21 fibroblasts did the ODC signal increase significantly with DFMO, but its presence was still greater in the T21 cells compared to D21 control cells (Figure 7A). APP could be seen throughout the cytoplasm of the fibroblast, with limited overlap with the nucleus, while ODC appeared both throughout the cytoplasm and in bright puncta (Figure 7B). Taking the data from individual cells for both APP and ODC, we compared them relative to one another between both cell lines and with DFMO treatments (Figure 7D). Linear regression of single-cell fluorescence for APP and ODC showed significant association between cell types and treatments, with an R^2^-value greater than 0.70 and p < 0.0001, indicating a strongly significant correlation.

**Figure 7:**
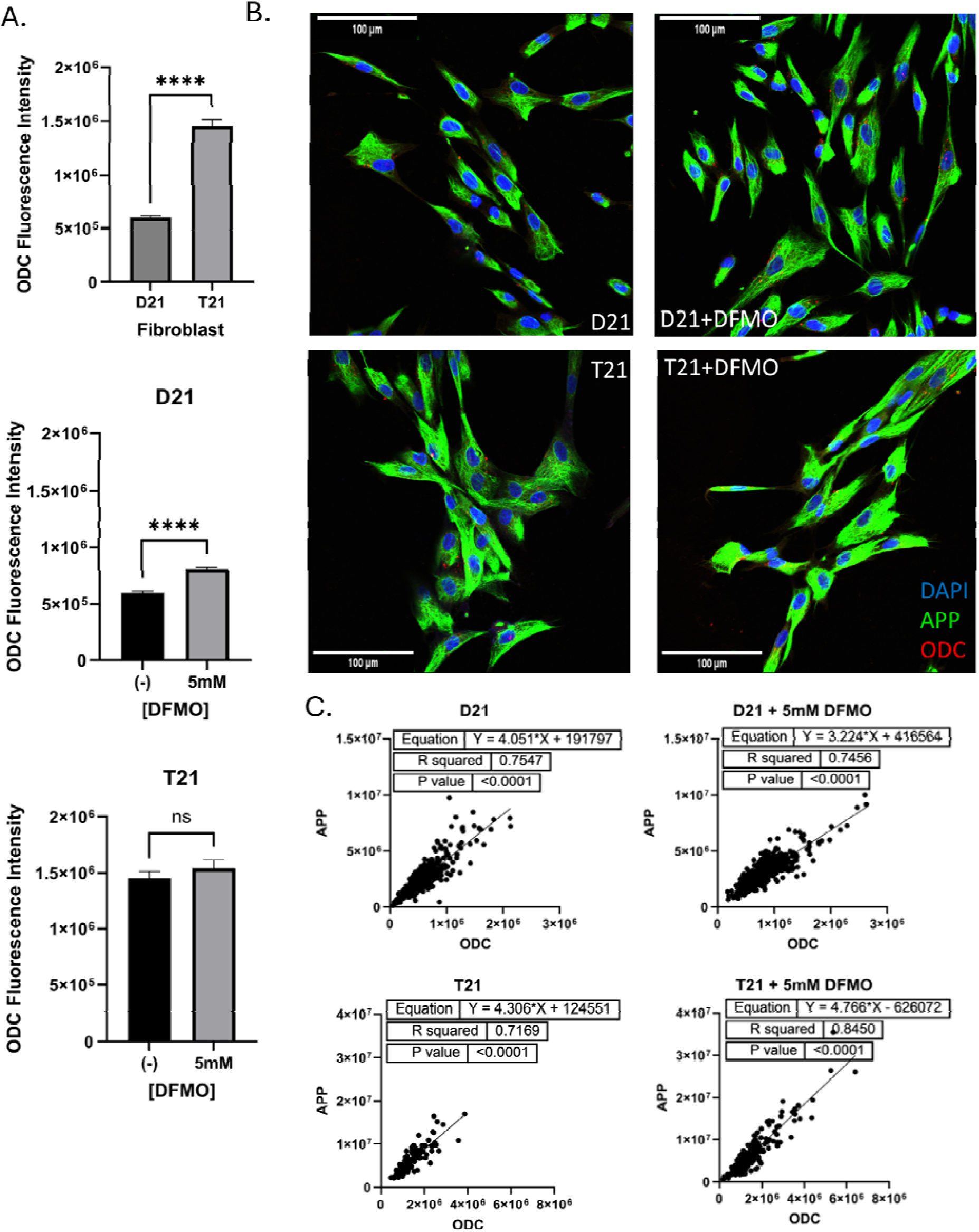
ODC Expression and its Correlation with APP in Normosomic and Trisomic Human Fibroblasts Treated with DFMO: A) Relative fluorescence of ODC as compared to each treated and untreated cell line with trisomic comparison to normosomic cells. B) Representative confocal images of human fibroblast cell cultures treated with DFMO. Results shown as mean SEM, n=200 (cells). **** indicates p < 0.0001, *** indicated p < 0.001, ** indicates p < 0.01, * indicates p < 0.05 as determined using two-way ANOVA with a post hoc Tukey’s test. C) Linear regression analysis of APP and ODC in normosomic and trisomic human fibroblasts treated with 5 mM DFMO.

### Protein expression and colocalization with A***β*** plaques in DS-AD brain tissue

Human brain tissues collected from patients with DS-AD, early-onset AD, and late-onset AD were lysed and analyzed with Western blots for the previously described proteins and compared to a HC case (See Supplemental Table 1 and Figure 8). Neither EOAD or LOAD differed from the control in Arginase 1, but the DS-AD brain showed twice the density (Figure 8A). EOAD had the greatest significant increase in ODC levels compared to the HC, with DS-AD following, but the levels of ODC in LOAD cases were not significantly different from the HC cases (Figure 8B). Spermine synthase levels were not significantly different from HC in the EOAD group, but the levels were significantly increased in the DS-AD group and to a lesser extent in the LOAD group (Figure 8C). Spermidine synthase levels were only significantly elevated in the LOAD brain compared to all other groups (Figure 8D). SAT1 relative levels were reduced with 50% in the DS-AD group, but not significantly different to the HC group in either the LOAD or EOAD groups (Figure 8E), suggesting a differential mechanism for PA dysregulation in DS cases compared to the general population. BACE1 relative levels were nearly six times higher in the DS-AD group compared to the HC group, while the BACE1 levels in the EOAD group exhibited were nearly twice as high as in the HC group (Figure 8F). The LOAD group did not exhibit a significant increase in BACE1 levels. These findings suggest differential mechanisms for the PA pathway dysregulation between LOAD, EOAD, and DS-AD groups, supporting continued investigation of this pathway in these conditions.

**Figure 8:**
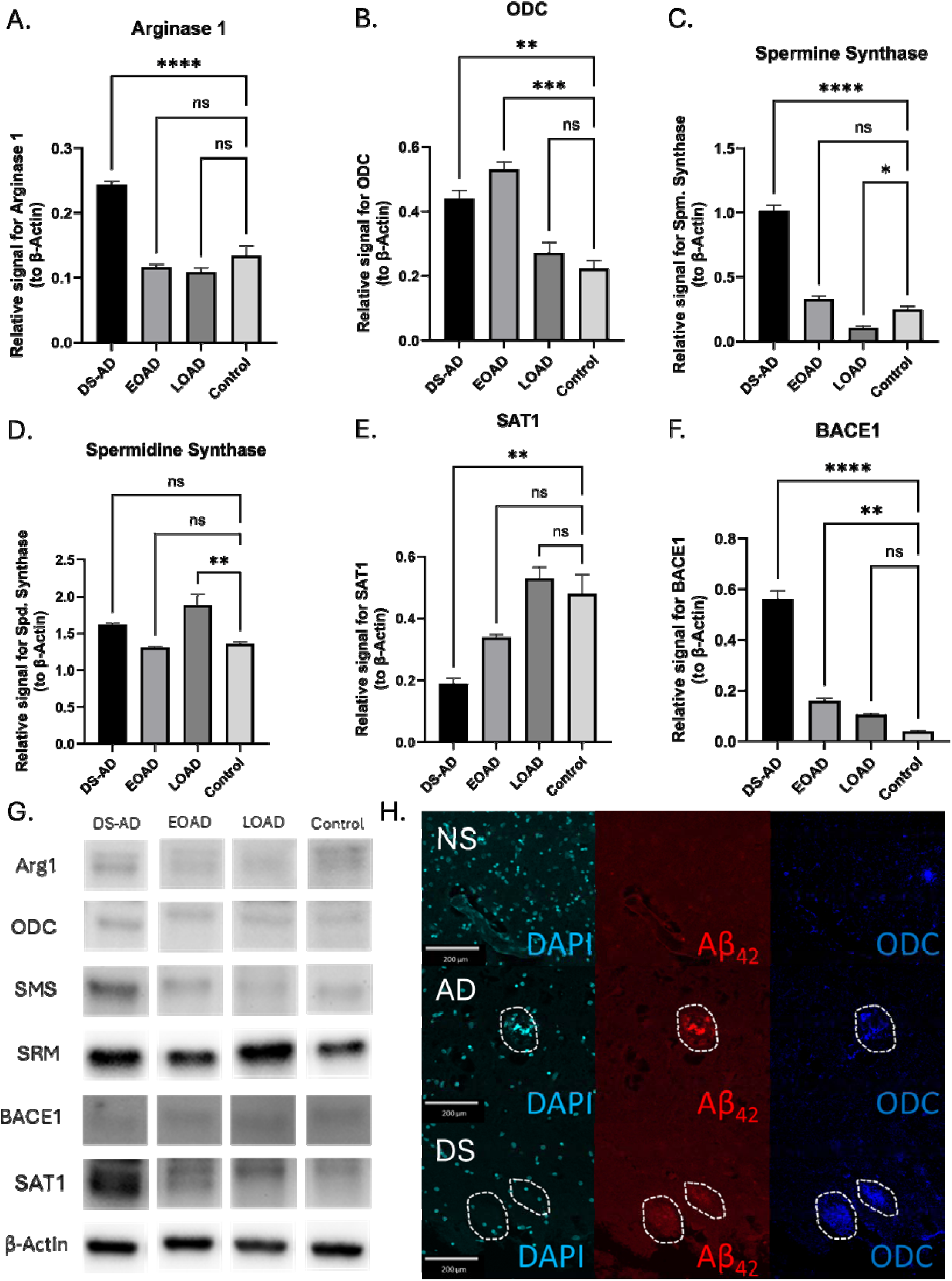
Altered PA Biosynthesis in DS-AD Brains: Protein Expression and Colocalization with Aβ Plaque: A) Western blot densitometry of human brain tissue lysate derived from DS-AD, EOAD, LOAD, and lung cancer (control) patients for A) arginase 1, B) ODC, C) spermine synthase, D) spermidine synthase, E) SAT1, and F) BACE1 with B-Actin as the control. G) Representative images of WB bands. H) Confocal images of a beta-amyloid plaque colocalizing with ODC in human brain tissue of normosomic, Alzheimer’s patient, and Down syndrome individuals with DAPI as a nuclear marker. Results shown as mean SEM, n=3. **** indicates p < 0.0001, *** indicated p < 0.001, ** indicates p < 0.01, * indicates p < 0.05 as determined using two-way ANOVA with a post hoc Tukey’s test.

Using immunofluorescence staining for beta-amyloid (Aβ42) and ODC (see Supplemental Table 1 for antibody details), we examined co-labeling of these two markers in the hippocampus of AD and DS (non-AD) individuals (Figure 8H). We found little presence of plaques in the normosomic (NS) brain, expectedly, and demonstrated few regions of increased ODC presence. Alternatively, both AD and DS saw plaques forming, and in these regions was a greater presence of ODC than the surrounding region. In the DS brain, ODC had other pockets of increased presence, but the plaques were not as distinct.

## DISCUSSION

In this study, we found that there was a significant increase in PA levels in both T21 cells and DS-AD or AD tissues compared to HC or D21 cell systems. Our findings demonstrated significantly elevated levels of all three major PAs - putrescine, spermidine, and spermine - in hippocampal tissue from AD patients compared to HC, and abnormal levels of PA synthesizing enzymes in the hippocampus of DS-AD, LOAD, EOAD cases compared to HC. This broad dysregulation of PA metabolism aligns with previous studies showing altered PA levels in AD brain tissue [56,57]. The concurrent elevation of spermidine and spermine levels suggests a comprehensive disruption of PA homeostasis rather than a pathway-specific alteration, and differences in dysregulation mechanisms between DS-AD and AD brains. The relatively small sample size (n=3), though yielding statistically significant results, underscores the need for validation in larger cohorts to confirm these preliminary findings. This study unveils a previously underexplored facet of AD and DS-AD pathophysiology, suggesting that the excessive accumulation of higher-order PAs could contribute to the underlying pathophysiology of AD, as evidenced by multiple in vitro models presented herein.

Our data suggest that Aβ increased levels of PAs in a dose dependent fashion, possibly via increased expression of ODC (Fig. 2). This upregulation of ODC could be driven by Aβ-induced cellular stress, as Aβ is known to activate pathways like the MAPK signaling cascade [56] and the unfolded protein response (UPR) [57], both of which are associated with the regulation of stress-responsive genes, including ODC. Consequently, this may represent an adaptive cellular attempt to buffer oxidative damage and mitigate the toxic effects of Aβ. Inhibiting ODC with the irreversible inhibitor DFMO decreased the deleterious effects of Aβ aggregation, oxidative stress, and in turn apoptosis in mouse hippocampal neurons. Together, these data provide the first evidence of the relationship between PAs and Aβ-induced toxicity and cell death, which has not been explored previously but is a vital step in understanding the complex mechanisms of dysfunction in the development of AD and may also lead to development of novel treatment avenues. Others have found direct and indirect evidence that elevated Aβ*_42_* levels can increase PA concentrations from analyses of rat embryonic hippocampal cells transfected with Aβ*_42_*, AD brain tissue, and hair samples from AD patients [27,30,33]. Our findings here expand previous findings, leading to the conclusion that PAs are upregulated by Aβ*_42_*in a dose dependent manner and thatthe amount of Aβ*_42_* aggregation is directly correlated to the intracellular concentration of PAs. Indeed, levels of PAs are reported to increase in AD brain tissue [33]. Although biogenic PAs can accelerate the rate of fibrillation of Aβ_42_ significantly [48], the expression of PAs in cell lines transfected with Aβ*_42_* had not been explored prior to this study. On the other hand, reduced putrescine and elevated spermine and spermidine levels in temporal cortex of patients with AD have previously been reported [56], suggesting a complex relationship between amyloid and PA metabolismThis dysregulation of the PA pathway could be involved not only in amyloid aggregation processes, but also in more fundamental and basic neuronal function such as calcium flux and glutamate homeostasis [58], and had not been examined in the DS-AD or DS brain prior to the present studies to our knowledge Previous work suggests that Aβ*_42_*aggregation upregulates PA levels by increasing expression of ODC, the rate-limiting enzyme of the PA biosynthesis pathway, responsible for the conversion of ornithine into putrescine [59,60]. In addition, ODC has previously been shown to exhibit increased activity in the presence of oxidative stress as well as translocation from the nucleus to the cytoplasm in AD brain tissue [36,43]. As expected, there was a significant increase in the ODC/actin ratio in cells exposed to increasing levels of Aβ*_42_* compared to cells exposed to IRES at the 24-hour timepoint. The ratio of ODC to beta-actin was greater in the Aβ*_42_* transfected lysates, allowing us to draw the conclusion that Aβ*_42_* increases PA levels by increased expression of ODC. The number of ODC immunoreactive neurons have been found to be increased in AD brains relative to control brains [29] further providing a link between Aβ_42_ toxicity and ODC expression and supporting our findings presented herein. While previous studies have shown that oxidative stress leads to greater ODC activity [36], the findings presented in Figure 2 are novel and warrant further investigation into the relationship between Aβ_42_ expression and ODC levels. This upregulation of PAs could be directly linked to the increase in ODC driven by Aβ-induced cellular stress response [56,57].

In addition to the effects of Aβ_42_ on PAs observed *in vitro*, Aβ_42_ aggregation kinetics were altered *in vitro* by the addition of putrescine (100 μM), strongly suggesting a direct effect of this PA precursor on the aggregation properties of Aβ_42_. It has been shown earlier that PAs are capable of promoting Aβ fibrillation via modulation of aggregation pathways and [47] suggested that this could be due to the fact that the three PAs share a similar binding mode to monomeric Aβ_42_peptide, leading to the acceleration of the aggregation process of Aβ_42_. Luo and collaborators suggested that this process was different for each PA, suggesting that very specific targeting molecules may have to be developed to successfully reduce amyloid aggregation via the PA pathway.

Having established that Aβ*_42_* aggregation increased PA levels in HT22 cells at least partially via increase of ODC, we subsequently investigated whether inhibition of PA synthesis by ODC inhibition would affect Aβ_42_ levels. DFMO has been proven to irreversibly inhibit ODC *in vitro* and *in vivo* [46,61]. Indeed, DFMO treatment of the HT22 cells transfected with Aβ*_42_* resulted in a significant decrease in PA levels in cells transfected with Aβ*_42_*. Further, inhibition of PA synthesis decreased Aβ*_42_* aggregation, reduced oxidative stress, and protected against apoptosis.

Others have demonstrated increased Aβ_42_ aggregation in human cell lines derived from people with DS [38] as well as Ts16 mouse-derived cell lines [62]. The Caviedes group demonstrated that Aβ_42_ overexpression in cortical Ts16 neurons was directly related to cell death [62], confirming the results observed herein. Further, when treated with DFMO, we found a reduction of amyloid fluorescence in the DS-Htk trisomy cell line – a cell that naturally over-expresses APP and amyloid [49,63]. Htk cells displayed a significant decrease in Aβ aggregation (Figure 5B). Further, upon treatment with DFMO, there was a significant decrease in PA levels in the Htk cells which was not significantly different from H1b control cells in both putrescine and spermine levels (Figure 5 C-E). Taken together, these data suggest that DFMO treatment could potentially reduce the aggregation of Aβ peptide in individuals with DS-AD, which could lead to a successful intervention strategy since DFMO has been proposed already as a treatment for other conditions, for example for neuroblastomas [64] or in diabetes to preserve β-cells [65].

As reported by others [66], BACE1 levels were significantly increased in DS-AD brain tissue, to a much greater extent than the increase seen in LOAD or EOAD brain tissues (Fig. 8). Since BACE1 expression levels are driven by APP levels [67], this is mostly likely because the APP gene is located on the Chr. 21 [6], and highly responsible for the early onset of AD pathology in DS. DFMO substantially reduced BACE1 expression in vitro, pointing to an interaction of the PA pathways also for cleavage of APP. According to our results, PA levels may modulate BACE1 expression through its transcription, potentially offering a mechanism through which PA imbalance could enhance amyloid-β production in DS-AD. This observation aligns with recent studies exploring the role of PAs in neurodegenerative processes, such as the work by Polis et al., 2021, which proposes AD as a chronic maladaptive PA stress response [60].

Significant differences were observed in terms of the PA synthetic enzymes between the DS-AD and LOAD or EOAD cases. The association between DS and AD is well documented [6] but the relationship between the polyamine pathway dysregulation and AD pathology in DS had not been investigated to date. Here we demonstrate alterations in DS brain tissue and fibroblasts related to PA expression and amyloid accumulation as well as the effects of PA inhibition on DS cell viability and protein homeostasis.

Although others have reported alterations of the PA pathway in AD and DS mouse models or cell systems [55], this is the first study to present evidence for involvement of this pathway in amyloid aggregation and cell viability. Moving forward, research should aim to establish the temporal relationship between PA accumulation and AD progression, clarifying the cellular mechanisms underlying the selective elevation of spermidine and spermine. Investigating these dysregulated PA levels across various brain regions and disease stages may yield insights into their role in AD pathogenesis and a potential development of DFMO for treatment options.

## DATA AVAILABILITY

All the datasets generated and analyzed during the current study are also available from the corresponding author. Source data is available for Figs. and Supplementary Figs. in the associated source data file. Source data are provided with this paper.

## ADDITIONAL INFORMATION

Correspondence and requests for materials should be addressed to D.P **Consent**: All brain tissues used for this study were obtained via a materials transfer agreement between DU and CU Anschutz. All cases included were obtained via a professional brain bank and with postmortem consent from the next of kin. The IRB approval and consent included storage of brain tissue and de-identified data in a brain bank and dissemination to other investigators.

## Acknowledgements

Supported by an R21 from the NIA R21AG070297, R01 AG061566-02, a grant from the Bright Focus Foundation (CA2018010), CONICYT for funding of Basal Centre, CeBiB, FB0001 and P09-022-F from ICM-ECONOMIA (Chile) and ICBM Enlaces at the University of Chile (570266).

## Conflict of interest statement

The authors declare no competing financial interests. Pablo Caviedes has IP protection on the H1B and HTk cell lines.

---

**Supplemental Table 1.** Antibodies utilized in immunohistochemistry (IHC) and Western blot (WB) experiments at indicated concentrations according to manufacturer’s recommendations for specific applications.

| Antibody | Manufacturer | Application | Concentration |
| --- | --- | --- | --- |
| Hoechst stain | Sigma (14530) | IHC | 1 ug/mL in PBS |
| MOAB-2 | Abcam (ab126649) | IHC | 5 ug/mL in primary solution |
| B-Amyloid peptide | Aves (ABN) | ICC | 1:2000 in blocking |
| APP A4 | EMD Millipore, MAB348 | IHC | 5 ug/mL in primary solution |
| ODC | Abcam (ab97395) | WB | 1:1000 in blocking |
| ODC | Abcam (193338) | IHC | 1:1000 in blocking |
| BACE1 | Abcam (183612) | WB | 1:1000 in blocking |
| Spermidine synthase | Abcam (241496) | WB | 1:1000 in blocking |
| Spermine synthase | Abcam (156879) | WB | 1:1000 in blocking |
| Arginase 1 | Cell Signaling (93668) | WB | 1:1000 in blocking |
| SAT1 | ThermoFisher (PA1-16992) | WB | 1:1000 in blocking |
| B-actin | Abcam (ab8227) | WB | 1:1000 in blocking |
| Anti-mouse HRP | BioRad (1706516) | WB | 1:500 in PBS-T |
| Anti-rabbit HRP | BioRad (1706515) | WB | 1:500 in PBS-T |

**Supplemental Table 1.**
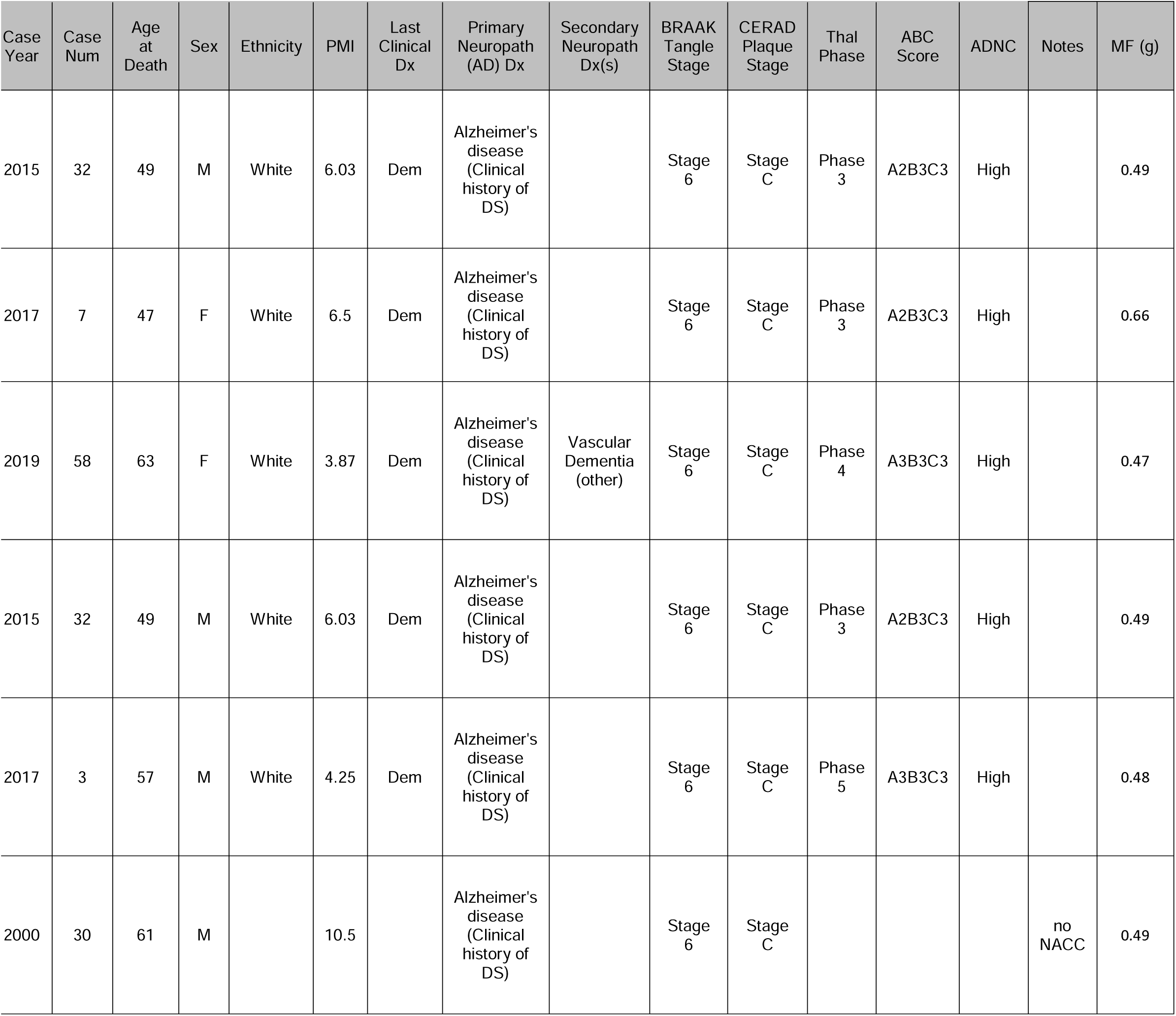
The hippocampal brain tissue used was obtained from a collaboration with Dr. Ann-Charlotte Granholm and the MUSC Carroll Campbell Jr. Neuropathology laboratory. This South Carolina Brain Bank contains frozen and fixed tissues from >250 Alzheimer cases and Controls, and all cases have undergone neuropathological and clinical staging including the ABC assessment as described by Jack et al [50] and recently amended by DeTure and Dickson [51] and Aldecoa et al., [10]

## Notes

### Summary of Updates

Revising the list of authors to match as they appear on the manuscript.

## Literature cited

[1] D. Hamlett E, A. Boger H, Ledreux A, M. Kelley C, J. Mufson E, F. Falangola M, et al. Cognitive Impairment, Neuroimaging, and Alzheimer Neuropathology in Mouse Models of Down Syndrome. Curr Alzheimer Res 2015;13:35–52. 10.2174/1567205012666150921095505.

[2] Hartley D, Blumenthal T, Carrillo M, DiPaolo G, Esralew L, Gardiner K, et al. Down syndrome and Alzheimer’s disease: Common pathways, common goals. Alzheimer’s Dement 2015;11:700–9. 10.1016/j.jalz.2014.10.007.

[3] Yates CM, Simpson J, Maloney AFJ, Gordon A, Reid AH. ALZHEIMER-LIKE CHOLINERGIC DEFICIENCY IN DOWN SYNDROME. Lancet 1980;316:979. 10.1016/S0140-6736(80)92137-6.

[4] Wiseman FK, Al-Janabi T, Hardy J, Karmiloff-Smith A, Nizetic D, Tybulewicz VLJ, et al. A genetic cause of Alzheimer disease: mechanistic insights from Down syndrome. Nat Rev Neurosci 2015;16:564–74. 10.1038/nrn3983.

[5] Wohlfert AJ, Phares J, Granholm A-C. The mTOR Pathway: A Common Link Between Alzheimer’s Disease and Down Syndrome. J Clin Med 2024;13. 10.3390/jcm13206183.

[6] Alldred MJ, Martini AC, Patterson D, Hendrix J, Granholm A-C. Aging with Down Syndrome-Where Are We Now and Where Are We Going? J Clin Med 2021;10. 10.3390/jcm10204687.

[7] Mann DMA, Esiri MM. The pattern of acquisition of plaques and tangles in the brains of patients under 50 years of age with Down’s syndrome. J Neurol Sci 1989;89:169–79. 10.1016/0022-510X(89)90019-1.

[8] Lemere CA, Blusztajn JK, Yamaguchi H, Wisniewski T, Saido TC, Selkoe DJ. Sequence of Deposition of Heterogeneous Amyloid β-Peptides and APO E in Down Syndrome: Implications for Initial Events in Amyloid Plaque Formation. Neurobiol Dis 1996;3:16–32. 10.1006/nbdi.1996.0003.

[9] Leverenz JB, Raskind MA. Early Amyloid Deposition in the Medial Temporal Lobe of Young Down Syndrome Patients: A Regional Quantitative Analysis. Exp Neurol 1998;150:296–304. 10.1006/exnr.1997.6777.

[10] Aldecoa I, Barroeta I, Carroll SL, Fortea J, Gilmore A, Ginsberg SD, et al. Down Syndrome Biobank Consortium: A perspective. Alzheimers Dement 2024;20:2262–72. 10.1002/alz.13692.

[11] Guo T, Zhang D, Zeng Y, Huang TY, Xu H, Zhao Y. Molecular and cellular mechanisms underlying the pathogenesis of Alzheimer’s disease. Mol Neurodegener 2020;15:40. 10.1186/s13024-020-00391-7.

[12] Billings LM, Oddo S, Green KN, McGaugh JL, LaFerla FM. Intraneuronal Aβ Causes the Onset of Early Alzheimer’s Disease-Related Cognitive Deficits in Transgenic Mice. Neuron 2005;45:675–88. 10.1016/j.neuron.2005.01.040.

[13] Hanon O, Vidal J, Lehmann S, Bombois S, Allinquant B, Tréluyer J, et al. Plasma amyloid levels within the Alzheimer’s process and correlations with central biomarkers. Alzheimer’s Dement 2018;14:858–68. 10.1016/j.jalz.2018.01.004.

[14] Chen JX, Yan S Du. Amyloid-β-Induced Mitochondrial Dysfunction. J Alzheimer’s Dis 2007;12:177–84. 10.3233/JAD-2007-12208.

[15] Bartley MG, Marquardt K, Kirchhof D, Wilkins HM, Patterson D, Linseman DA. Overexpression of Amyloid-β Protein Precursor Induces Mitochondrial Oxidative Stress and Activates the Intrinsic Apoptotic Cascade. J Alzheimer’s Dis 2012;28:855–68. 10.3233/JAD-2011-111172.

[16] Cheignon C, Tomas M, Bonnefont-Rousselot D, Faller P, Hureau C, Collin F. Oxidative stress and the amyloid beta peptide in Alzheimer’s disease. Redox Biol 2018;14:450–64. 10.1016/j.redox.2017.10.014.

[17] Childs AC, Mehta DJ, Gerner EW. Polyamine-dependent gene expression. Cell Mol Life Sci 2003;60:1394–406. 10.1007/s00018-003-2332-4.

[18] Kanemura A, Yoshikawa Y, Fukuda W, Tsumoto K, Kenmotsu T, Yoshikawa K. Opposite effect of polyamines on In vitro gene expression: Enhancement at low concentrations but inhibition at high concentrations. PLoS One 2018;13:e0193595. 10.1371/journal.pone.0193595.

[19] Heby O. Role of Polyamines in the Control of Cell Proliferation and Differentiation. Differentiation 1981;19:1–20. 10.1111/j.1432-0436.1981.tb01123.x.

[20] Weiger TM, Hermann A. Cell proliferation, potassium channels, polyamines and their interactions: a mini review. Amino Acids 2014;46:681–8. 10.1007/s00726-013-1536-7.

[21] Eisenberg T, Knauer H, Schauer A, Büttner S, Ruckenstuhl C, Carmona-Gutierrez D, et al. Induction of autophagy by spermidine promotes longevity. Nat Cell Biol 2009;11:1305– 14. 10.1038/ncb1975.

[22] Zhang H, Alsaleh G, Feltham J, Sun Y, Napolitano G, Riffelmacher T, et al. Polyamines Control eIF5A Hypusination, TFEB Translation, and Autophagy to Reverse B Cell Senescence. Mol Cell 2019;76:110–125.e9. 10.1016/j.molcel.2019.08.005.

[23] Zhang H, Simon AK. Polyamines reverse immune senescence via the translational control of autophagy. Autophagy 2020;16:181–2. 10.1080/15548627.2019.1687967.

[24] Chattopadhyay MK, Tabor CW, Tabor H. Polyamines Are Not Required for Aerobic Growth of *Escherichia coli*L: Preparation of a Strain with Deletions in All of the Genes for Polyamine Biosynthesis. J Bacteriol 2009;191:5549–52. 10.1128/JB.00381-09.

[25] Minois N, Carmona-Gutierrez D, Madeo F. Polyamines in aging and disease. Aging (Albany NY) 2011;3:716–32. 10.18632/aging.100361.

[26] Soulet D, Rivest S. Polyamines play a critical role in the control of the innate immune response in the mouse central nervous system. J Cell Biol 2003;162:257–68. 10.1083/jcb.200301097.

[27] Choi MH, Kim K, Kim IS, Lho DS, Chung BC. Increased hair polyamine levels in patients with alzheimer’s disease. Ann Neurol 2001;50:128–128. 10.1002/ana.1086.

[28] Morrison LD, Bergeron C, Kish SJ. Brain S-adenosylmethionine decarboxylase activity is increased in Alzheimer’s disease. Neurosci Lett 1993;154:141–4. 10.1016/0304-3940(93)90191-M.

[29] Bernstein H-G, Müller M. Increased immunostaining for l-ornithine decarboxylase occurs in neocortical neurons of Alzheimer’s disease patients. Neurosci Lett 1995;186:123–6. 10.1016/0304-3940(95)11301-C.

[30] Yatin SM, Yatin M, Aulick T, Ain KB, Butterfield DA. Alzheimer’s amyloid β-peptide associated free radicals increase rat embryonic neuronal polyamine uptake and ornithine decarboxylase activity: protective effect of vitamin E. Neurosci Lett 1999;263:17–20. 10.1016/S0304-3940(99)00101-9.

[31] Yatin SM, Yatin M, Varadarajan S, Ain KB, Butterfield DA. Role of spermine in amyloidL?-peptide-associated free radical-induced neurotoxicity. J Neurosci Res 2001;63:395–401. 10.1002/1097-4547(20010301)63:5<395::AID-JNR1034>3.0.CO;2-Q.

[32] Seidl R, Beninati S, Cairns N, Singewald N, Risser D, Bavan H, et al. Polyamines in frontal cortex of patients with Down syndrome and Alzheimer disease. Neurosci Lett 1996;206:193–5. 10.1016/S0304-3940(96)12451-4.

[33] Inoue K, Tsutsui H, Akatsu H, Hashizume Y, Matsukawa N, Yamamoto T, et al. Metabolic profiling of Alzheimer’s disease brains. Sci Rep 2013;3:2364. 10.1038/srep02364.

[34] Gilad GM, Gilad VH. Novel polyamine derivatives as neuroprotective agents. J Pharmacol Exp Ther 1999;291:39–43.

[35] Gilad GM, Gilad VH, Wyatt RJ. Accumulation of exogenous polyamines in gerbil brain after ischemia. Mol Chem Neuropathol 1993;18:197–210. 10.1007/BF03160034.

[36] Saito K, Packianathan S, Longo LD. Free radical-induced elevation of ornithine decarboxylase activity in developing rat brain slices. Brain Res 1997;763:232–8. 10.1016/S0006-8993(97)00414-9.

[37] Murray Stewart T, Dunston TT, Woster PM, Casero RA. Polyamine catabolism and oxidative damage. J Biol Chem 2018;293:18736–45. 10.1074/jbc.TM118.003337.

[38] Pegg AE. Regulation of Ornithine Decarboxylase. J Biol Chem 2006;281:14529–32. 10.1074/jbc.R500031200.

[39] Perez-Leal O, Merali S. Regulation of polyamine metabolism by translational control. Amino Acids 2012;42:611–7. 10.1007/s00726-011-1036-6.

[40] Uemura T, Nakamura M, Sakamoto A, Suzuki T, Dohmae N, Terui Y, et al. Decrease in acrolein toxicity based on the decline of polyamine oxidases. Int J Biochem Cell Biol 2016;79:151–7. 10.1016/j.biocel.2016.08.039.

[41] Yoshida M, Tomitori H, Machi Y, Hagihara M, Higashi K, Goda H, et al. Acrolein toxicity: Comparison with reactive oxygen species. Biochem Biophys Res Commun 2009;378:313–8. 10.1016/j.bbrc.2008.11.054.

[42] Moghe A, Ghare S, Lamoreau B, Mohammad M, Barve S, McClain C, et al. Molecular Mechanisms of Acrolein Toxicity: Relevance to Human Disease. Toxicol Sci 2015;143:242–55. 10.1093/toxsci/kfu233.

[43] Nilsson T, Bogdanovic N, Volkman I, Winblad B, Folkesson R, Benedikz E. Altered subcellular localization of ornithine decarboxylase in Alzheimer’s disease brain. Biochem Biophys Res Commun 2006;344:640–6. 10.1016/j.bbrc.2006.03.191.

[44] Skatchkov SN, Woodbury-Fariña MA, Eaton M. The Role of Glia in Stress. Psychiatr Clin North Am 2014;37:653–78. 10.1016/j.psc.2014.08.008.

[45] Nakanishi S, Cleveland JL. Polyamine Homeostasis in Development and Disease. Med Sci (Basel, Switzerland) 2021;9. 10.3390/medsci9020028.

[46] Gomes GM, Dalmolin GD, Bär J, Karpova A, Mello CF, Kreutz MR, et al. Inhibition of the Polyamine System Counteracts β-Amyloid Peptide-Induced Memory Impairment in Mice: Involvement of Extrasynaptic NMDA Receptors. PLoS One 2014;9:e99184. 10.1371/journal.pone.0099184.

[47] Luo J, Yu C-H, Yu H, Borstnar R, Kamerlin SCL, Gräslund A, et al. Cellular Polyamines Promote Amyloid-Beta (Aβ) Peptide Fibrillation and Modulate the Aggregation Pathways. ACS Chem Neurosci 2013;4:454–62. 10.1021/cn300170x.

[48] Kabir A, Jash C, Payghan P V., Ghoshal N, Kumar GS. Polyamines and its analogue modulates amyloid fibrillation in lysozyme: A comparative investigation. Biochim Biophys Acta - Gen Subj 2020;1864:129557. 10.1016/j.bbagen.2020.129557.

[49] Cárdenas A. Cell Lines Derived from Hippocampal Neurons of the Normal and Trisomy 16 Mouse Fetus (a Model for Down Syndrome) Exhibit Neuronal Markers, Cholinergic Function, and Functional Neurotransmitter Receptors. Exp Neurol 2002;177:159–70. 10.1006/exnr.2002.7957.

[50] Jack CR, Knopman DS, Weigand SD, Wiste HJ, Vemuri P, Lowe V, et al. An operational approach to National Institute on Aging–Alzheimer’s Association criteria for preclinical Alzheimer disease. Ann Neurol 2012;71:765–75. 10.1002/ana.22628.

[51] DeTure MA, Dickson DW. The neuropathological diagnosis of Alzheimer’s disease. Mol Neurodegener 2019;14:32. 10.1186/s13024-019-0333-5.

[52] Betancourt L, Rada P, Hernandez L, Araujo H, Ceballos GA, Hernandez LE, et al. Micellar electrokinetic chromatography with laser induced fluorescence detection shows increase of putrescine in erythrocytes of Parkinson’s disease patients. J Chromatogr B 2018;1081–1082:51–7. 10.1016/j.jchromb.2018.02.015.

[53] Hernandez L, Joshi N, Murzi E, Verdeguer P, Mifsud JC, Guzman N. Collinear laser-induced fluorescence detector for capillary electrophoresis. J Chromatogr A 1993;652:399–405. 10.1016/0021-9673(93)83259-U.

[54] Dada OO, Huge BJ, Dovichi NJ. Simplified sheath flow cuvette design for ultrasensitive laser induced fluorescence detection in capillary electrophoresis. Analyst 2012;137:3099–101. 10.1039/c2an35321k.

[55] Páez X, Rada P, Hernández L. Neutral amino acids monitoring in phenylketonuric plasma microdialysates using micellar electrokinetic chromatography and laser-induced fluorescence detection. J Chromatogr B Biomed Sci Appl 2000;739:247–54. 10.1016/S0378-4347(99)00536-8.

[56] Palavicini JP, Wang C, Chen L, Hosang K, Wang J, Tomiyama T, et al. Oligomeric amyloid-beta induces MAPK-mediated activation of brain cytosolic and calcium-independent phospholipase A2 in a spatial-specific manner. Acta Neuropathol Commun 2017;5:56. 10.1186/s40478-017-0460-6.

[57] Lindholm D, Korhonen L, Eriksson O, Kõks S. Recent Insights into the Role of Unfolded Protein Response in ER Stress in Health and Disease. Front Cell Dev Biol 2017;5. 10.3389/fcell.2017.00048.

[58] Bowie D, Lange GD, Mayer ML. Activity-Dependent Modulation of Glutamate Receptors by Polyamines. J Neurosci 1998;18:8175–85. 10.1523/JNEUROSCI.18-20-08175.1998.

[59] Lane DJR, Bae D-H, Siafakas AR, Suryo Rahmanto Y, Al-Akra L, Jansson PJ, et al. Coupling of the polyamine and iron metabolism pathways in the regulation of proliferation: Mechanistic links to alterations in key polyamine biosynthetic and catabolic enzymes. Biochim Biophys Acta - Mol Basis Dis 2018;1864:2793–813. 10.1016/j.bbadis.2018.05.007.

[60] Polis B, Karasik D, Samson AO. Alzheimer’s disease as a chronic maladaptive polyamine stress response. Aging (Albany NY) 2021;13:10770–95. 10.18632/aging.202928.

[61] Nie L, Feng W, Diaz R, Gratton MA, Doyle KJ, Yamoah EN. Functional Consequences of Polyamine Synthesis Inhibition by l-α-Difluoromethylornithine (DFMO). J Biol Chem 2005;280:15097–102. 10.1074/jbc.M409856200.

[62] Arriagada C, Bustamante M, Atwater I, Rojas E, Caviedes R, Caviedes P. Apoptosis is directly related to intracellular amyloid accumulation in a cell line derived from the cerebral cortex of a trisomy 16 mouse, an animal model of Down syndrome. Neurosci Lett 2010;470:81–5. 10.1016/j.neulet.2009.12.062.

[63] Cárdenas AM, Ardiles AO, Barraza N, Baéz-Matus X, Caviedes P. Role of Tau Protein in Neuronal Damage in Alzheimer’s Disease and Down Syndrome. Arch Med Res 2012;43:645–54. 10.1016/j.arcmed.2012.10.012.

[64] Tangella AV, Gajre AS, Chirumamilla PC, Rathhan P V. Difluoromethylornithine (DFMO) and Neuroblastoma: A Review. Cureus 2023;15:e37680. 10.7759/cureus.37680.

[65] Sims EK, Kulkarni A, Hull A, Woerner SE, Cabrera S, Mastrandrea LD, et al. Inhibition of polyamine biosynthesis preserves β cell function in type 1 diabetes. Cell Reports Med 2023;4:101261. 10.1016/j.xcrm.2023.101261.

[66] Miners JS, Morris S, Love S, Kehoe PG. Accumulation of insoluble amyloid-β in down’s syndrome is associated with increased BACE-1 and neprilysin activities. J Alzheimers Dis 2011;23:101–8. 10.3233/JAD-2010-101395.

[67] Luo G, Xu H, Huang Y, Mo D, Song L, Jia B, et al. Deposition of BACE-1 Protein in the Brains of APP/PS1 Double Transgenic Mice. Biomed Res Int 2016;2016:8380618. 10.1155/2016/8380618.

